# Spike and host glycan determinants of HKU1 airway tropism

**DOI:** 10.64898/2026.08.05.741156

**Authors:** Robert Creutznacher, Louisa E. Wallace, Oliver J. Debski-Antoniak, Ieva Drulyte, Ruben J. G. Hulswit, Jeffrey Beekman, Frank J. M. van Kuppeveld, Berend-Jan Bosch, Cornelis A.M. de Haan, Martin Frank, Daniel L. Hurdiss, Raoul J. de Groot

## Abstract

Human coronavirus HKU1 comprises two distinct serotypes, A and B, whose spike proteins are substantially divergent. Here, we show that spikes from both serotypes preferentially bind 9-*O*-acetylated α2,8-linked disialosides. Cryo-electron microscopy of the B-type N5 spike reveals a conserved extended binding site in domain S1^A^ that accommodates both the terminal and penultimate sialic acid residues, with interactions involving the penultimate residue substantially enhancing binding. Spike N-glycan processing modulates affinity and linkage selectivity; glycans flanking the binding pocket offer a plausible structural basis for these effects. In contrast to the HKU1-A spike, which adopts open S1^B^-up conformations upon ligand binding, the N5 apo structure showed that the ligand-binding site was already formed and the e1 relay element register-shifted in most protomers. Nevertheless, we detected neither spontaneous opening nor a transition to an S1^B^-up state following ligand binding. These findings, obtained with a minimally modified ectodomain, differ from recent reports of ligand-independent opening. Molecular dynamics simulations indicated that membrane-embedded GT3, but not GD3, presents its glycan chain in a geometry compatible with S1^A^-mediated engagement. Concordantly, in human nasal epithelial cultures cell surface GT3-like *O*-acetylated trisialoside glycotopes were detected in ciliated cells, linking their cell-type-specific presentation to HKU1 tropism.

## INTRODUCTION

Humans are subject to recurrent upper respiratory tract infections caused by two closely related common cold betacoronaviruses (CoVs) belonging to the subgenus *Embecovirus.* These viruses, called HKU1 and OC43, originated from rodent reservoirs and entered the human population independently, either directly or, as in the case of OC43, via a succession of intermediate hosts. OC43 emerged relatively recently, apparently from a bovine coronavirus (BCoV) spillover an estimated 70-120 years ago with the most recent common OC43 ancestor dated to the 1950s^1^. HKU1 presumably entered the human population decades if not centuries earlier, although its date of origin cannot be estimated^2^. Its immediate ancestor remains to be identified and there is a paucity of sequence data for HKU1 field variants sampled before 2006. Circulation under continuous immune selection has driven viral divergence through antigenic drift and recombination, giving rise to three HKU1 genotypes, designated A through C. Moreover, two serotypes, A and B, can be distinguished that differ particularly in their spike proteins (S), with genotype C encompassing recombinant viruses carrying B-type S genes in a genotype A background^3^. As an indication for evolutionary distance, the A- and B-type S proteins are only 84% identical. For reference, BCoV and OC43 S proteins are far more closely related, sharing more than 91% sequence identity.

HKU1 and OC43 differ from other human CoVs in that they use cell surface glycans decorated with terminal 9-*O*-acetylated sialic acid as essential primary receptors^4,5^. Binding to these ligands occurs through a generally conserved binding site in N-terminal domain S1^A^ of the spike protein S (for S domain organization see Fig. S1). Another envelope protein, the hemagglutinin esterase (HE) - among CoVs, unique to embecoviruses - serves as a receptor-destroying enzyme by virtue of its sialate-9-*O*-acetylesterase activity^6,7^. S and HE are thought to function as a two-component system for dynamic receptor interaction during (pre)attachment, promoting enzyme-driven virion motility to traverse the mucus layer and glycocalyx and to allow receptor-assisted diffusion along the cell surface^8,9^. Upon S1^A^-mediated cell surface attachment to sialoglycan-based primary receptors, HKU1 S binds a secondary, proteinaceous receptor - transmembrane serine protease 2 (TMPRSS2) - via domain S1^B10,11,12^. Exposure of the TMPRSS2-binding site requires conformational changes, apparently conditional on S1^A^-sialoside association^13^.

Whereas cellular expression of TMPRSS2 is a prerequisite for HKU1 S-mediated fusion and viral entry, ligand preference and receptor promiscuity of the S1^A^ domain may be a contributing if not a key factor for host-, organ- and cell tropism. Capitalizing on our knowledge of CoV HE substrate specificities, we previously generated a library of 9-mono-*O*- and 7,9-di-*O*-acetylated sialosides through chemo-enzymatic synthesis^14^. Screening of this library by glycan array analysis with the S protein of serotype A HKU1 strain Caen1 as analyte revealed a strong selectivity for α2,8-linked 9-*O*-acetyldisialoside motifs. Questions remain, however, whether the results obtained for HKU1 field variant Caen1 reflect its full spectrum of ligand usage and whether they are representative of that of HKU1 variants in general. Here we provide experimental evidence that selective binding to disialoside-based receptors is not a serotype- or strain-specific trait. We show that the sialoglycan binding profile of the type B spike of HKU1 strain N5 is highly similar to that of A-type Caen1 S, and, if anything, even more selective for α2,8-linked 9-*O-*Ac-Sias. Single-particle cryo-EM analysis of the B-type S protein, corroborated by molecular dynamics and structure-guided mutagenesis, found the extended disialoside-specific binding site recently defined for A-type HKU1 N1 spike^15^ to be conserved despite considerable sequence variation in surrounding areas. The findings provide detailed insight into how the penultimate, reducing-end Sia (Sia1) of the disialoside glycotope is accommodated within this site and how protein-Sia1 contacts contribute to binding affinity and receptor selection. However, whereas the cryo-EM data give a protein-structural explanation for the preferential binding of α2,8-linked disialosides, we show that N-glycans shielding the terminal Sia2-binding site not only modulate binding affinity but contribute to ligand selectivity.

In A-type spikes, disialoside-binding selects for and/or stabilizes local and long-range conformational changes in S1^A^, which are an essential prelude to S1^B^ transitioning from a down (‘closed’) to an up (’open’) state^13,15^. In contrast, but consistent with an earlier study^16^, we find that in our B-type S *apo* structure, the ‘activating’ conformational changes in S1^A^ are already present in more than 80% of the protomers. Despite this, the trimer did not open spontaneously, differing from more recent reports^17,18^. Moreover, even upon disialoside binding, when virtually all S1^A^ domains adopt the activated state, the HKU1-B spikes remain fully closed.

To study the importance of S1^A^-mediated primary receptor binding during natural infection, we used air-liquid interface cultures as a model for human nasal respiratory epithelia. Reflecting HKU1 cell tropism^19^, S1A probes bound preferentially to ciliated cells and colocalized with A2B5-reactive, lipid-extractable glycoconjugates consistent with GT3-like glycolipids. Given that glycan-dependent attachment precedes TMPRSS2-dependent membrane fusion^10^, expression of cell surface 9-*O*-Ac disialoside motifs may be a key determinant of HKU1 host cell tropism.

## RESULTS

### Glycan Binding Profiles of HKU1 Serotypes A and B Revealed by Biolayer Interferometry

A previous screening of a library of biotinylated *O*-acetylated sialoglycans by glycan microarray analysis with probes comprising viral receptor-binding lectin domains expressed as Fc fusion proteins^14^ revealed a strong selectivity of the serotype A HKU1 strain Caen1 S1^A^ domain for α2,8-linked 9-*O*-Ac-disialosides. This particular ligand preference over α2,3- and 2,6-linked monosialosides was recently confirmed for HKU1 N1^15^, a closely related A-type strain differing from Caen1 in S1^A^ by only four residues (I32V, K84Y, I166K and D272N). To study whether the results obtained for the serotype A strains reflected the full spectrum of HKU1 ligand usage and whether the observed ligand preference is representative of that of HKU1 variants in general rather than a serotype-specific trait, we compared the glycan binding properties of Caen1 and serotype B strain N5 S proteins by biolayer interferometry (BLI). In contrast to glycan microarray analysis and beyond a qualitative assessment, BLI allows label-free real-time determination of association and dissociation kinetics. Biotinylated α2,3-, α2,6-, and α2,8-linked mono-(9-*O*-Ac) and di-*O*-acetylated (7,9-*O*-Ac_2_) sialoglycans were immobilized on biosensors. As analytes we used trimeric S ectodomains with S1^A^ domains presented in the native structural context to ensure that the observed binding profiles reflect those of the native S proteins. Both HKU1-A and -B S trimers preferentially bound to 9-*O*-Ac-Sia-α2,8-Sia with the 9-*O*-Ac group being essential, corroborating the earlier observations made for the HKU1-A S1^A^-Fc fusion protein (Fig. 1b). The binding responses of the A- and B-type S proteins were comparable, with B-type S consistently showing stronger binding.

**Fig. 1.**
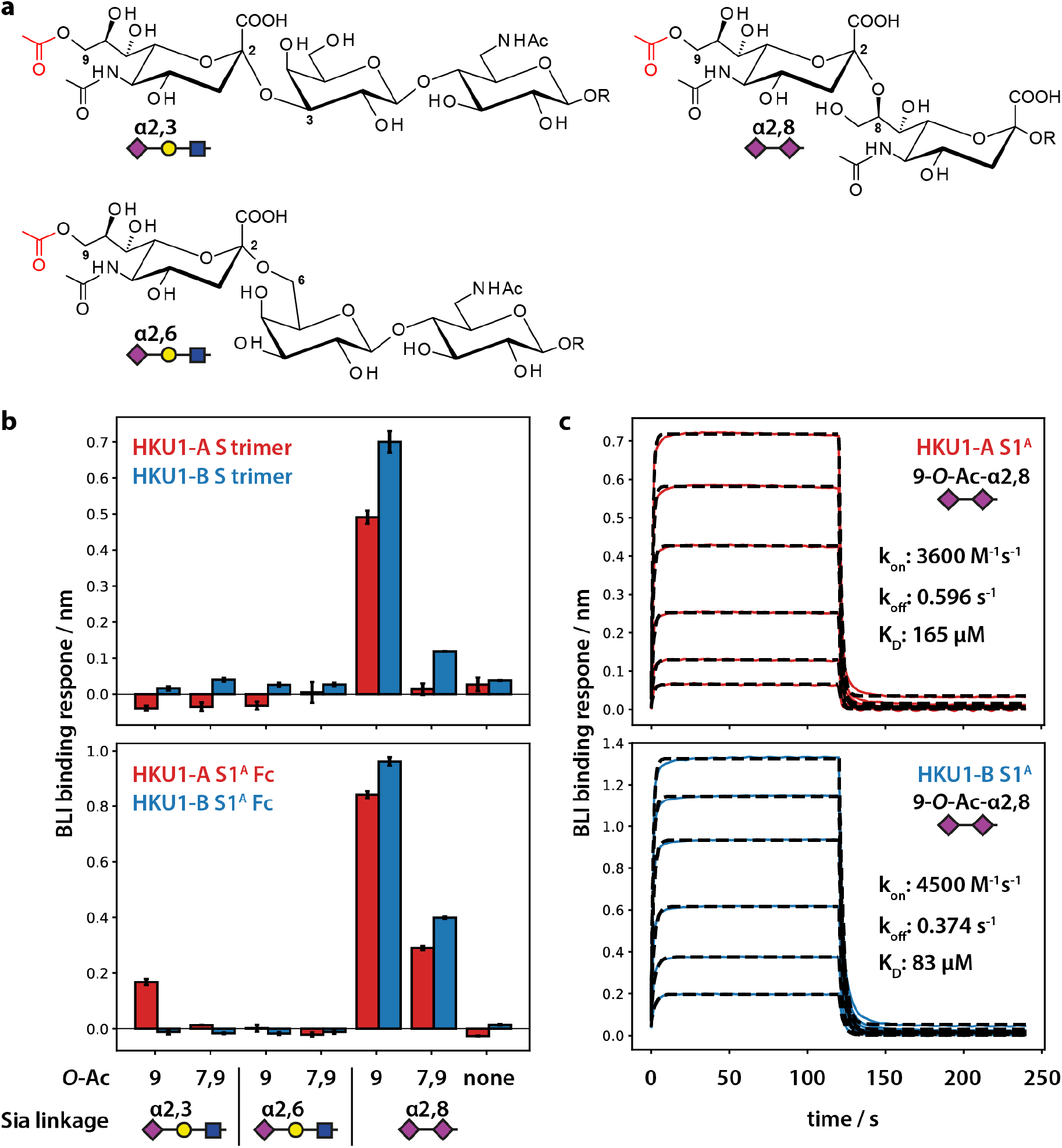
Sialoglycan binding of HKU1 S proteins. **a.** Structures of different synthetic 9-*O*-Ac sialosides (R = Lc-biotin^14^). **b.** 9-*O*-acetylated and 7,9-*O*-diacetylated sialoglycans were tested for binding of trimeric HKU1 S ectodomains (0.38 µM) and S1^A^-Fc proteins (2.5 µM) in a BLI-based glycan array. Error bars denote standard deviations between duplicate measurements from pristine sensors. **c.** Binding curves of HKU1-A and -B S1^A^ monomers interacting with 9-O-Ac-Sia-α2,8-Sia at different protein concentrations (6.25–95 µM) were fit to a kinetic 1:1 binding model (dashed lines). The derived dissociation constants K_D_ agree with results from steady state analysis of the same data (Fig. S2).

Low expression yields of S ectodomains precluded BLI experiments at higher analyte concentrations. Consequently, low-affinity interactions with 9-*O*-Ac-sialosides in other glycosidic linkages may have gone undetected. We therefore repeated the experiments at higher concentrations using bivalent S1^A^-Fc fusion proteins as probes. Again, both the A- and B-type analytes preferentially bound to 9-*O*-Ac-Sia-α2,8-disialosides and, to a lesser extent, to 7,9-di-*O*-Ac_2_-Sia-α2,8-Sia. However, under these conditions, S1^A^ of HKU1-A, but not of HKU1-B, showed low affinity binding also to 9-*O*-Ac-α2,3-sialyllactose. To measure kinetic binding parameters, BLI analyses were performed with S1^A^ monomers, allowing for 1:1 binding and avoiding confounding effects of multivalency (Fig. 1c). Binding to 9-*O*-Ac-Sia-α2,8-Sia was analyzed at a range of protein concentrations, yielding binding curves that quickly reached equilibrium binding levels. Association rates, estimated from curve fitting to a kinetic binding model, were between 3000 and 4500 M^-1^ s^-^1. They were thus 5-100 times higher than those determined for human influenza A virus (IAV) HAs^20^, but still far below the diffusion limit, suggesting limited accessibility of the S1^A^ binding site. Dissociation rate constants were in the same range as those of the IAV HAs, 0.6 s^-1^ for HKU1-A and 0.4 s^-1^ for HKU1-B S1^A^, corresponding to residence times of approximately 1.7 s and 2.5 s, respectively. Dissociation constants (K_D_) averaged from duplicate experiments were in the order of 170 µM and 110 µM for HKU1-A and HKU1-B, respectively, corresponding to those estimated by independent curve fitting of steady-state binding levels (150 and 60 µM, respectively, Fig. S2). The K_D_ values determined by us are considerably higher, up to 30-fold, than those recently reported by others^15^ but may better approximate the actual binding affinities. Whereas our probes carry complex and hybrid *N*-glycans, those used by Jin et al. were expressed in *N*-acetylglucosaminyltransferase I deficient cells. Replacement of natural N-glycans by immature oligomannose-type Man_5_-GlcNAc_2_ chains strongly increases binding affinity, as shown previously^21^ and below.

The combined findings show that the S protein of HKU1-B strain N5, like those of HKU1-A strains Caen1 and N1, displays a strong preference for α2,8-linked 9-*O*-Ac-disialoside-based receptors. The observation of variants of either serotype preferentially binding to disialoside motifs supports the hypothesis that this is an adaptation for replication in the human respiratory tract^14^. While the Caen1 and N5 S proteins displayed comparable binding kinetics towards the preferred ligand, N5 was the more selective of the two.

### Cryo-EM analysis of HKU1-B strain N5 spike complexes

To gain insight into the structural basis for HKU1 S ligand preference and receptor-induced conformational changes, we collected single-particle cryo-EM data sets of the S ectodomain of HKU1 strain N5 both in the unbound state and in complex with the 9-*O*-Ac-disialoside. In contrast to recent studies, we used a minimally modified expression product with only the furin cleavage site inactivated but, to avoid potential consequences for local protein conformational integrity and stability, without the Pro substitutions introduced by others^15,17,18,22^. Reconstructions yielded density maps of 2.6 and 2.4 Å global resolution, respectively (Fig. 2a, Table S1, Figs. S3–S6).

**Fig. 2.**
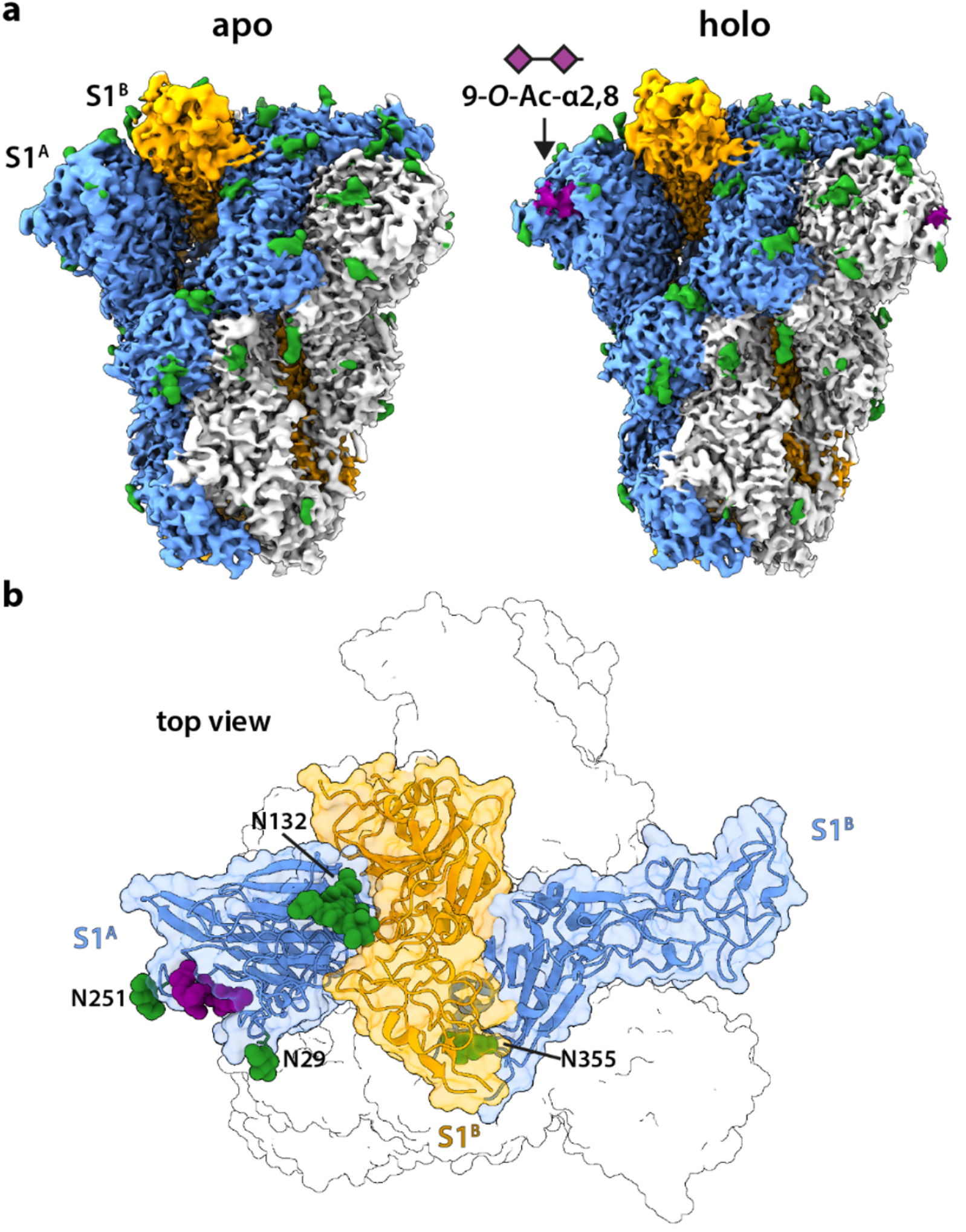
Closed structure of the HKU1-B S protein. **a**. EM density maps of the trimeric HKU1-B S ectodomain in the absence and presence of a 9-*O*-Ac disialoside ligand (purple). Individual protomers are colored blue, yellow, or white. Densities of modeled *N*-glycans are indicated in green. **b**. Top view of the modelled complex between S and its sialoglycan ligand (surface representation in white). The structures of one representative S1^B^ domain (yellow), wedged between domains S1^A^ and S1^B^ of a neighboring protomer (blue), are highlighted in ribbon representation. The disialoside ligand and select N-glycans are shown in purple and green, respectively.

An unbound, endoglycosidase H-treated S structure of HKU1-B N5 was determined previously at a resolution of 4 Å^16^ as one of the first CoV S structures reported. In comparison, we were able to model an additional 247 amino acids and 30 carbohydrates per protomer with an overall Cα RMSD of 1.1 Å. This allowed us to visualize the membrane-proximal regions and large parts of the S1^B^ domain, which harbors the binding site for HKU1’s proteinaceous secondary receptor TMPRSS2^11,12^.

The HKU1-B N5 S *apo*- and *holo* structures are highly similar to each other (RMSD 0.34 Å), both resembling the non-complexed N5 structure described by Kirchdoerfer et al.^16^. They adopt a virtually identical, closed prefusion conformation with the S1^B^ domains in the down conformation. Our N5 *apo* S structure differs in several respects from that described for A-type HKU1 spikes. First, the tip of the HKU1-B S1^B^ domain is shifted by 9 Å, such that the TMPRSS2 binding site, shielded in HKU1-A S in its closed state, appears more exposed. When the S1^B^ domain is modelled to bind TMPRSS2 in this topology, most of the clashes with the adjacent S1 protomer, noted by others^11^, are resolved (Fig. S7). Another difference concerns the S1^A^ conformation and ligand-induced conformational changes. In non-complexed A-type spikes, the N-terminal segment (res. 29-37), designated the e1 element, is predominantly disordered^13,15^. Upon ligand binding, whether by induced fit or conformational selection, the e1 element assumes a stable topology such that the 9-*O*-Ac-Sia binding site becomes fully structured. This is associated with long-range changes involving conserved e1 residues R34 and S36 and a simultaneous breaking of two interstrand backbone hydrogen bonds (S36–D76 and Y38–F74) and their re-formation with new partners (R34–D76 and S36–F74) in a ‘register shift’. These local conformational changes result in an inward rotation of the top of the S1^A^ domain (termed subdomain S1^A1^) relative to the S1^A^ base (subdomain S1^A2^) with downstream consequences for S1^A^-S1^B^ interactions.

Overall, our N5 *apo* structure, like the one determined by Kirchdoerfer et al., more closely resembles HKU1-A Caen1 S in its disialoside-complexed closed, down state (Cα RMSD 2.7 Å)^13^ and the closed, complexed, and non-complexed activated states recently described for HKU1-A N1 (RMSD 1.8 Å)^15^. A focused conformational analysis of the S1^A^ domain using 3D variability analysis^23^ estimated that approximately 87% of N5 protomers contained a formed terminal-Sia2 binding site and a register-shifted e1 element (Fig. S8). In the remaining protomers, the S1^A^ domains adopt an outwardly displaced conformation relative to the consensus map, consistent with the non-activated *apo* state described for HKU1-A S^13^.

In stark contrast to observations for A-type spikes, which showed the closed, *apo* state of S converting into open S1^B^-up conformations upon ligand binding, we detected only the closed, all-down conformation for complexed HKU1-B N5 spike. With ligand density visible at all symmetry-related sites in the reconstruction, i.e. at virtually complete ligand occupancy with all S protomers now in the S1^A^ activated state, the HKU1-B spike remained closed. The datasets were obtained under conditions identical to those in our prior study on HKU1-A Caen1 S^13^ with incubation of the S-disialoside complex for 10 min at RT. Even after extending the incubation time to 1 h and with the temperature elevated to 37 °C, the spike ectodomains remained fully closed (Fig. S9). Exhaustive 3D variability analysis did not reveal any indication of the presence of spike trimers in an open conformation (Fig. S8).

Our findings also differ from those of recent studies of the S proteins of HKU1-B strains N5 and N2. Whereas the *apo* structure of N2 S is similar to that of non-activated type A HKU1 spikes^18^, Wang et al.^17^ modelled the *apo* closed N5 structure as a uniformly pseudo-symmetric complex with each of the monomers differing in S1^A^ conformation. However, inspection of the corresponding density map suggested that also in their apo closed structure all e1 conformations are consistent with the non-activated state (Fig. S10). Even more puzzlingly, both groups reported spontaneous, ligand-independent transitions to S1^B^-up conformations. We note, however, that in both studies, the S ectodomains contained stabilizing 2P substitutions^22^, which have been described to affect S1^B^ dynamics in the case of the SARS-CoV-2 S^24^. Whereas the conflicting observations may stem from methodological differences (see Discussion), from a pragmatic perspective, the homogeneity of our data obtained for ligand-complexed HKU1-B N5 did allow us to gain high-resolution insight into the structural details of S1^A^-mediated sialoglycan binding.

### Structural Basis of HKU1-B Spike Interaction with Sialic Acid Ligands

Local refinement of S1^A^ accounted for minor conformational dynamics around the S1^A^-S1^B^ interface and allowed us to resolve the sialoglycan ligand (Fig. 3a, b). The terminal 9-*O*-Ac sialic acid (Sia2) is located within a canyon formed by e1 residues 29-31 together with residues 79-86, and by element e2 (residues 245–252). The binding site closely resembles those in other embecoviruses^21^ with all key contact residues conserved (Fig. S12). Among these, Asn26, Lys80, and Ser82 interact with terminal Sia2 directly, while Trp89 and Thr31 are essential for the formation of the pockets p1 and p2 that accommodate the 9-*O*- and 5-*N*-acetyl groups, respectively. In HKU1-B, Tyr28 flanks the glycerol side chain and forms an additional hydrogen bond with Sia2-O7. The Y28A substitution abrogates detectable binding to 9-*O*-Ac-disialosides in BLI analyses (Fig. 3c). Ordered water molecules apparently stabilize the ligand by bridging the Sia2-O3 and −5-*N*-Ac group to the surrounding protein through residues Ile235 and Thr31, respectively. The positioning of Sia2 and surrounding water molecules as deduced from the experimental map was supported by independent molecular dynamics (MD) simulations of the solvated glycan binding site (Fig. S13). Of note, in our structure Sia2 would adopt the natural chair conformation^25^, different from that reported by Wang et al.^17^, which was modelled to take on a highly unusual high-energy boat conformation with distorted bond angles (Fig. S14).

**Fig. 3.**
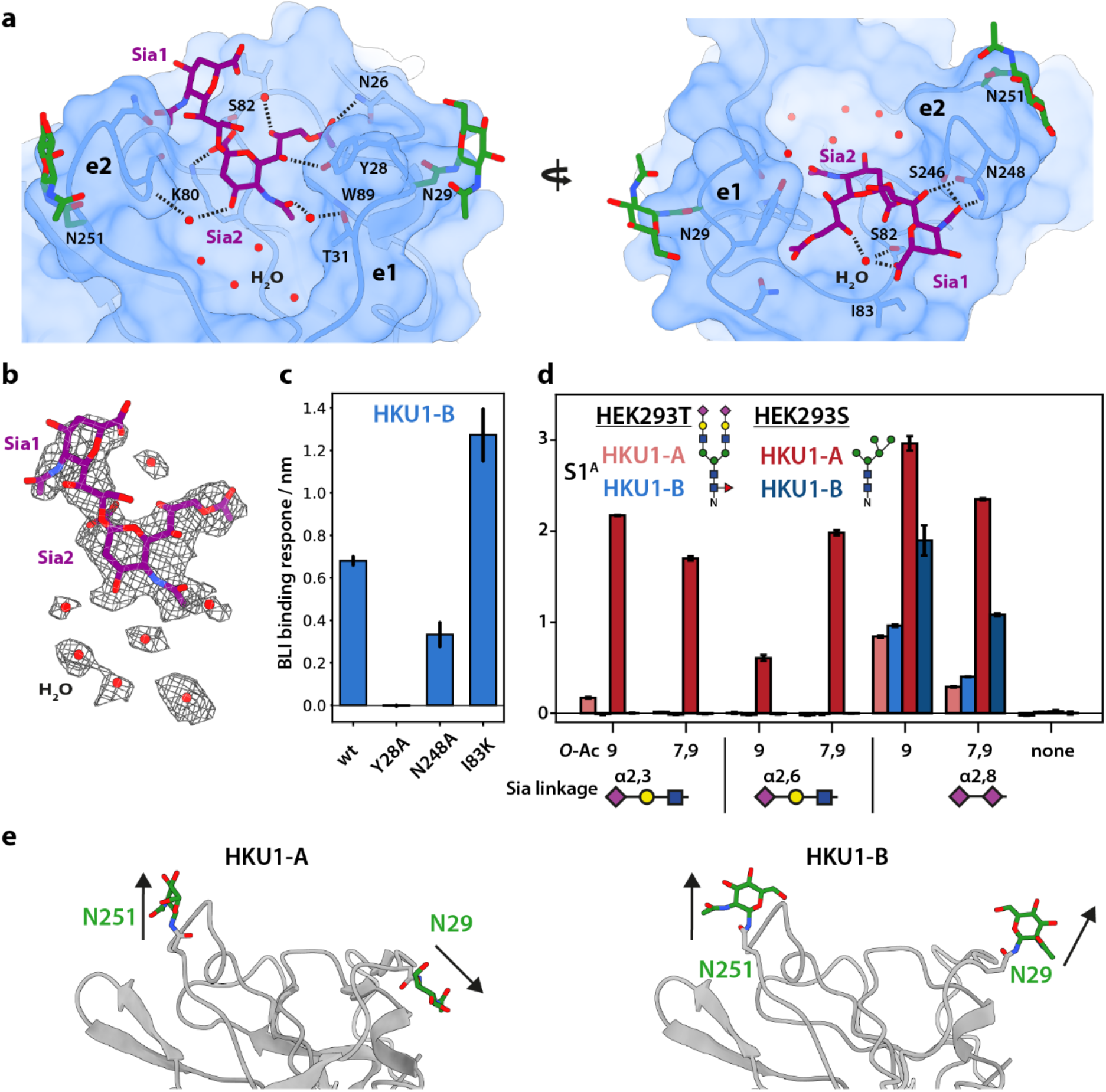
The binding site for 9-*O*-Ac α2,8-linked sialoglycans of HKU1-B S. **a.** The binding pose of the disialoside ligand (purple) is shown in a surface model of the S1^A^ domain. Hydrogen bonds with the protein and surrounding ordered water molecules are indicated. The binding site is flanked by the two N-glycosylated loops e1 and e2. **b.** Experimental EM density map of the ligand and surrounding water molecules. **c.** Binding of HKU1-B S1^A^-hFc point mutants to 9-*O*-Ac-Sia-α2,8-Sia as determined by BLI compared to that of the parental (wt) protein. Error bars denote standard deviations between duplicate measurements from pristine sensors. **d.** N-glycans modulate accessibility and selectivity of the sialoglycan binding site. Exclusively Man_5_ N-glycosylated HKU1 S1^A^–Fc proteins, produced in HEK293S GnTI(-) cells (bars in dark red and dark blue), show stronger binding than those with more extensively processed HEK293T-derived N-glycans produced in HEK293T cells (bars in light red and light blue). Error bars denote standard deviations between duplicate measurements from pristine sensors. WT data are reproduced from Fig. 1 for comparison. **e.** Binding patterns of HKU1-A and -B S glycoforms might be explained by the different orientation of N-glycans surrounding the binding site.

In our previous HKU1-A S structure, conformational heterogeneity and pronounced flexibility in the sialoglycan binding pocket limited resolution, obscuring any stabilizing contacts between the protein and the penultimate, reducing-end Sia1. Very recently^15^, the high-resolution structure of A-type strain N1 showed the existence of an extended binding site also accommodating the penultimate Sia residue (Sia1) (Fig. S15a). The current data provide structural evidence that this disialoside binding site is conserved in B-type S protein despite considerable sequence variation and the evolutionary distance separating A- and B-type spikes. The structure details how Sia1 interactions contribute to ligand binding by HKU1-B S. The cryo-EM map of the S-disialoside complex, as compared to that of the *apo* protein, revealed a substantial increase in e2 density as evidenced by a decrease of average Cα B-factors in e2 from 105 Å² to 63 Å². Apparently, this element becomes more ordered upon binding of the ligand, allowing for the formation of a third pocket p3 that can harbor the Sia1 5-*N*-Ac group, with options for hydrogen bonding of the 5-*N*-acetyl group with Ser246 and Asn248, and of Sia1 O7 with Asn248. N248A substitution reduced BLI binding levels to approximately half compared to wild-type protein (Fig. 3c, Fig. S12a). We also noted that as a natural polymorphism some HKU1-B field variants feature a Lys at position 83 (Fig. S12b), its side chain seemingly poised to form an additional salt bridge with the Sia1 carboxylate^13^. Predictably, I83K substitution resulted in a gain of function, substantially increasing disialoside binding, mostly due to an increase in the association rate (Fig. S15b).

To gain further insight into why α2,8-linked 9-*O*-Ac-Sia is favored over α2,3- and α2,6-linked sialosides, we performed MD simulations of HKU1-A and -B complexed with different sialoglycans (Fig. S16). The results suggest that the HKU1 S1^A^ domain can accommodate a terminal 9-*O*-Ac-Sia in more than one glycosidic context when the ligand is encountered in solution. However, in the case of the α2,6- and α2,3-linked sialoglycans, the modelled interactions involved predominantly the terminal 9-*O*-Ac-Sia2 without significant contributions from the 2^nd^ and 3^rd^ residues in the extended glycan chains. In contrast, the conformational flexibility conferred by α2,8-linkage - the adjacent sugar rings are separated by one or two additional C-bonds, respectively - allows 9-*O*-Ac-disialosides to dock into an extended binding site with Sia1 engaging in glycan-protein interactions at the base of e2 corresponding to those implied by cryo-EM.

### Spike N-glycan processing modulates binding affinity and ligand specificity

The cryo-EM data provide a protein-structural explanation for the preferential binding of HKU1-B N5 S proteins to α2,8-linked disialoside motifs over α2,6- and α2,3-linked monosialylated glycans. However, our previous observations for the S protein of HKU1-A Caen1 indicated a potential role also for two conserved N-glycans linked to Asn29 and Asn251, respectively, decorating the e1- and e2-loops and shielding the Sia1-binding site^21^. In apparent consequence, HKU1-A S1^A^-Fc fusion proteins, expressed in HEK293S GnTI-deficient cells to replace complex and hybrid *N*-glycans by immature oligomannose-type Man_5_-GlcNAc_2_ (M5) chains^26^, displayed enhanced binding to cell surface sialosides as measured by hemagglutination assay^21^. Furthermore, in the case of HKU1-B N5 S1^A^, M5-replacement of N-linked glycans reportedly resulted in a broader receptor binding profile when compared to that of its HEK293T-derived counterpart carrying more extensively processed N-glycans, at least when tested by glycan array analysis with probes multivalently presented on nanoparticles^27^. To further our understanding of how S N-glycosylation affects binding affinity and ligand selectivity, we performed BLI analysis with M5-glycosylated dimeric S1^A^-Fc fusion proteins (Fig. S17), testing them against our library of 9-*O*-Ac-sialosides. For HKU1-B strain N5 S1^A^-Fc, we observed a strongly increased binding (Fig. 3d) as compared to the protein produced in standard HEK293T cells, but selectivity for 9-*O*-Ac α2,8-linked disialosides was virtually maintained with no detectable binding to α2,3- and α2,6-linked sialosides. HKU1-A M5-S1^A^–Fc displayed enhanced binding to disialosides even beyond the levels measured for N5 S1^A^–Fc. However, in stark contrast to the N5 probe, M5-glycosylated Caen1 S1^A^ readily bound also to α2,3- and α2,6-linked 9-*O*-Ac-Sias, amplifying binding events that were barely (α2,3) or not at all (α2,6) detectable with the HEK293T-derived glycoform of the protein. The increased binding levels to these ligands relative to those to α2,8-linked disialosides were consistent over a range of lower probe concentrations (Fig. S18), indicating that they did not result from a general increase in affinity, but rather from a loss of linkage specificity. Apparently, mature glycans, attached to the S1^A^ e1 Asn29 and e2 Asn251 - elsewhere identified as bi- and tri-/tetra-antennary in HKU1-B S, respectively^28^ - not only lower binding affinity by reducing binding site availability and thereby the association rate^21^ but also contribute to ligand selectivity. The differential effect of M5 substitution, observed for HKU1-A and -B S, may be explained by differences in e1 conformations to the effect that the core N-acetylglucosamine moiety linked to Asn29 would extend from the respective proteins at different angles (Fig. 3e). Whereas in HKU1-A S the attached N-glycan would point away, it adopts a topology perpendicular to the plane of the binding site in HKU1-B S, such that an M5 high-mannose chain might still hamper the efficient docking of α2,3- and α2,6-linked sialoside-based ligands.

### Ciliated cells in human nasal epithelium display glycan-based receptors of HKU1

The analyses described so far were performed with synthetic glycans either in solution or in solid-phase BLI binding assays. To study HKU1 S-mediated attachment to natural cell surface sialoglycans, we used air-liquid interface cultures of fully differentiated human nasal epithelial cells (HNEC) as an in vitro model system mimicking the cell composition of the upper respiratory tract^29^. In accordance with previous findings^19^, our HNEC cultures supported HKU1 propagation as shown by immunofluorescence assay (IFA) with an HKU1-cross-reactive anti-mouse hepatitis virus polyclonal serum as well as with a monoclonal antibody (mAb) against double-stranded RNA as a marker for infection. Infection, observed at 24 h p.i., was restricted to ciliated cells, identified as such with β-tubulin as an intracellular marker (Fig. 4a). Confirming the key role of cell surface 9-*O*-Ac-Sias in HKU1 attachment^5^, infection was prevented by sialate-9-*O*-acetylesterase pretreatment of HNEC (Fig. 4b).

**Fig. 4.**
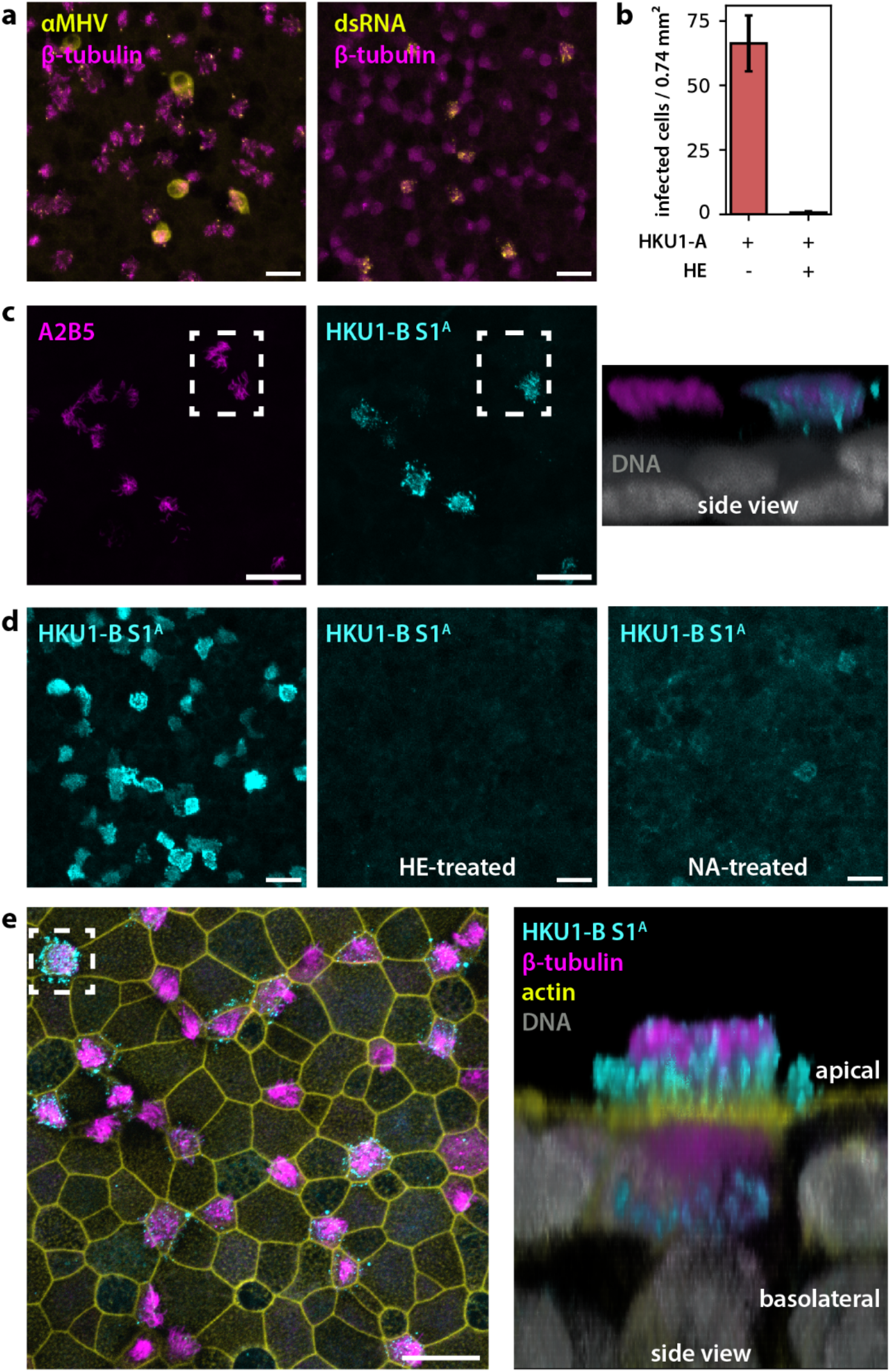
HKU1 host cell tropism assessed in human nasal epithelial cell cultures. **a.** HKU1-A Caen1 specifically targets ciliated cells. Infected cells were identified by IFA staining with a cross-reactive anti-MHV polyclonal serum or with a mAb against double-stranded RNA (yellow); β-tubulin served as a marker for ciliated cells (magenta). **b.** HKU1-A infects ciliated cells in a 9-*O*-Ac-Sia-dependent manner. The number of infected cells 24 h p.i. with and without pre-treatment with bovine CoV HE was determined based on confocal imaging. Error bars denote the standard deviation across three overview images from the two respective membrane inserts. **c.** Surface immunofluorescence staining with HKU1-B S1^A^ (cyan) colocalizing with the A2B5 glycotope in a subset of A2B5^+^ cells (magenta). **d.** HKU1-B S1^A^ specifically binds to cell surface 9-*O*-acetylated sialosides. Binding is lost upon prior treatment with bovine CoV HE or neuraminidase (NA). **e.** Sialoglycan receptors recognized by HKU1 S1^A^ (cyan) are exclusively present on a subset of ciliated cells. IFA co-staining for β-tubulin (magenta) and cell boundaries (actin, yellow); nuclei stained with DAPI (grey). The right panel shows a side view of the indicated area with HKU1 S1^A^ staining cilia on the apical side. Control experiments under mock conditions are shown in Fig. S19. Top view images in a, c, d and e are shown as maximum intensity projections of confocal stacks. Scale bars denote 20 µm.

Sialic acids, including those modified by 9-*O*-acetylation, generally occur as α2,3- and α2,6-linked terminal residues of the *N*- and *O*-linked glycan chains of cell surface proteins. In human lung tissue, terminal α2,8-linked disialosides are also found in *O*-glycans but have been detected less frequently^30^. They seem to occur more commonly on B- and C-series gangliosides like GD3 and GT3^31,32^. In a previous study, we showed that such gangliosides can function as primary receptors for HKU1^14^. Induction of their synthesis by over-expression of ST8SIA1 in HEK293T cells resulted in cell surface binding of HKU1 S and promoted infection of HKU1 S-pseudotyped viruses. Prompted by these findings, we studied the occurrence of gangliosides in formaldehyde-fixed, non-permeabilized HNEC by IFA. Staining for GD3 or 9-*O*-Ac-GD3 with mAbs R24^31^ and UM4D4^33^, respectively, produced no detectable staining, but a monoclonal antibody A2B5^34^ directed against trisialoside motifs typical for GT3-like gangliosides gave a bright surface staining restricted to ciliated cells (Fig. S19). Fluorescent areas appeared as isolated protrusions suggestive of cilia, extending approximately 10 µm above the nuclear layer on the apical side. Delipidation of HNEC with organic solvents prevented A2B5 staining, consistent with its binding of cell surface glycolipids (Fig. S20). Control experiments with *Sambucus nigra* agglutinin (SNA) lectin, primarily recognizing 2,6-linked sialoglycoproteins^35^, showed persistent albeit diminished apical surface staining of all cell types. The apparent presence of trisialoside glycolipids in human airway tissue is consistent with an early study in which GT3 was identified as a ganglioside of human lung^32^.

To study the distribution of HKU1 glycan-based ligands in comparison, non-permeabilized HNECs were subjected to IFA with S1^A^-Fc fusion proteins (Fig. 4c). In accordance with HKU1 host cell tropism, the 9-*O*-Ac-Sia-dependent binding of the HKU1-A and -B probes was restricted to ciliated cells (Fig. 4d, e). HKU1-B S1^A^ gave a brighter fluorescence signal than that of HKU1-A (Fig. S21), in line with the difference in their affinities towards 9-*O*-Ac-Sia-α2,8-Sia. Binding was lost upon sialate-de-*O*-acetylation or desialylation of cell surface glycans, providing evidence for binding specificity. While S1^A^-Fc fluorescence intensity varied between cells, it invariably coincided with A2B5 staining. However, a subset of A2B5+ ciliated cells showed no detectable binding of HKU1 S1^A^ (Fig. 4c). Of note, A2B5 binds both *O*-acetylated and non-*O*-acetylated GT3-type glycotopes^34^. The observations may thus be attributed to cell-to-cell diversity in levels of sialate-*O*-acetylation as was noted previously both in cultured cell populations as well as in natural tissues^36^. In direct competition assays, HKU1-B S1^A^ suppressed A2B5 binding in a concentration-dependent fashion, consistent with overlapping recognition of *O*-acetylated A2B5-reactive glycotopes (Fig. S22). Conversely, S1^A^ staining was detectable up to the highest tested concentration (0.2 mg/ml) of A2B5, perhaps indicating substantial differences in multivalent binding strength between the two probes or the existence of additional HKU1-binding glycotopes, e.g., disialoside motifs in *O*-linked glycans, beyond those recognized by A2B5.

Consistent with a substantial glycolipid contribution to HKU1 S1^A^ binding, cell surface staining was mostly lost upon HNEC delipidation (Fig. S20). Remarkably, in HNECs thus permeabilized, IFA with S1^A^-Fc fusion proteins resulted in a bright, perinuclear staining in all cell types, in accordance with our previous observations in deparaffinized sections of the human respiratory tract^14^. Apparently, large intracellular reservoirs of 9-*O*-Ac-Sia-sialosides as detectable by HKU1 S do not necessarily correlate with detectable cell surface presentation. Our combined findings suggest that cell surface A2B5-reactive, lipid-extractable GT3-like glycoconjugates may serve as primary HKU1 receptors. This would subsequently allow binding to the secondary proteinaceous receptor TMPRSS2 which, like the A2B5-type HKU1 S1^A^ ligands, preferentially occurs in ciliated cells.

### Disialoside accessibility in membrane-embedded gangliosides

S-mediated virion attachment to cell surface gangliosides is complicated by the proximity of the plasma membrane, the accessibility of ganglioside sialoglycans, the shielding of the S sialoside-binding site by N-linked glycans and even the positioning of the ligand-binding pocket, not at the tip but rather at the lateral side of the S headgroup. MD simulations of membrane-embedded, 9-*O*-acetylated gangliosides with increasing chain length showed the ceramide-linked Glc residue to be buried between the phospholipid head groups with the terminal Sia typically protruding from the outer leaflet by 6, 8 and 12 Å for GM3, GD3 and GT3, respectively (Fig. 5a). Attachment this close to the membrane might be hampered by the S-associated N-glycans, acting as spacer elements. To study this, we performed 100 ns MD simulations of a complex between a glycosylated HKU1-B S trimer and a 9-*O*-Ac GT3 ganglioside embedded in a phospholipid bilayer (Fig. 5b, c). The results show that binding to glycolipid ligands in a membrane environment is possible without clashes with the lipid bilayer but would indeed require substantial reorientation of the S1^A^ *N*-glycans at Asn29 and Asn251. In order to bind to GT3, spikes would have to approach the host cell membrane at an angle of about 32° (Fig. 5b), potentially reliant on stalk flexibility in the viral membrane^37,38^. Additional MD simulations that enforced complex formation between the S1^A^ domain and different gangliosides showed that binding to GM3 and GD3 resulted in substantial deviations from the average unbound ganglioside conformations. For GM3 these discrepancies seem too severe to be compatible with binding, whereas for GD3 they would at least incur a binding penalty. Thus, its unfavorable glycan presentation might make GD3 less suitable as a receptor. In contrast, the length of the GT3 pentasaccharide chain extending from the membrane in its average unbound conformation was geometrically compatible with binding of the terminal disialoside motif, accommodating both Sia2 and Sia1 in their respective binding sites similar to the free-state S-disialoside complex.

**Fig. 5.**
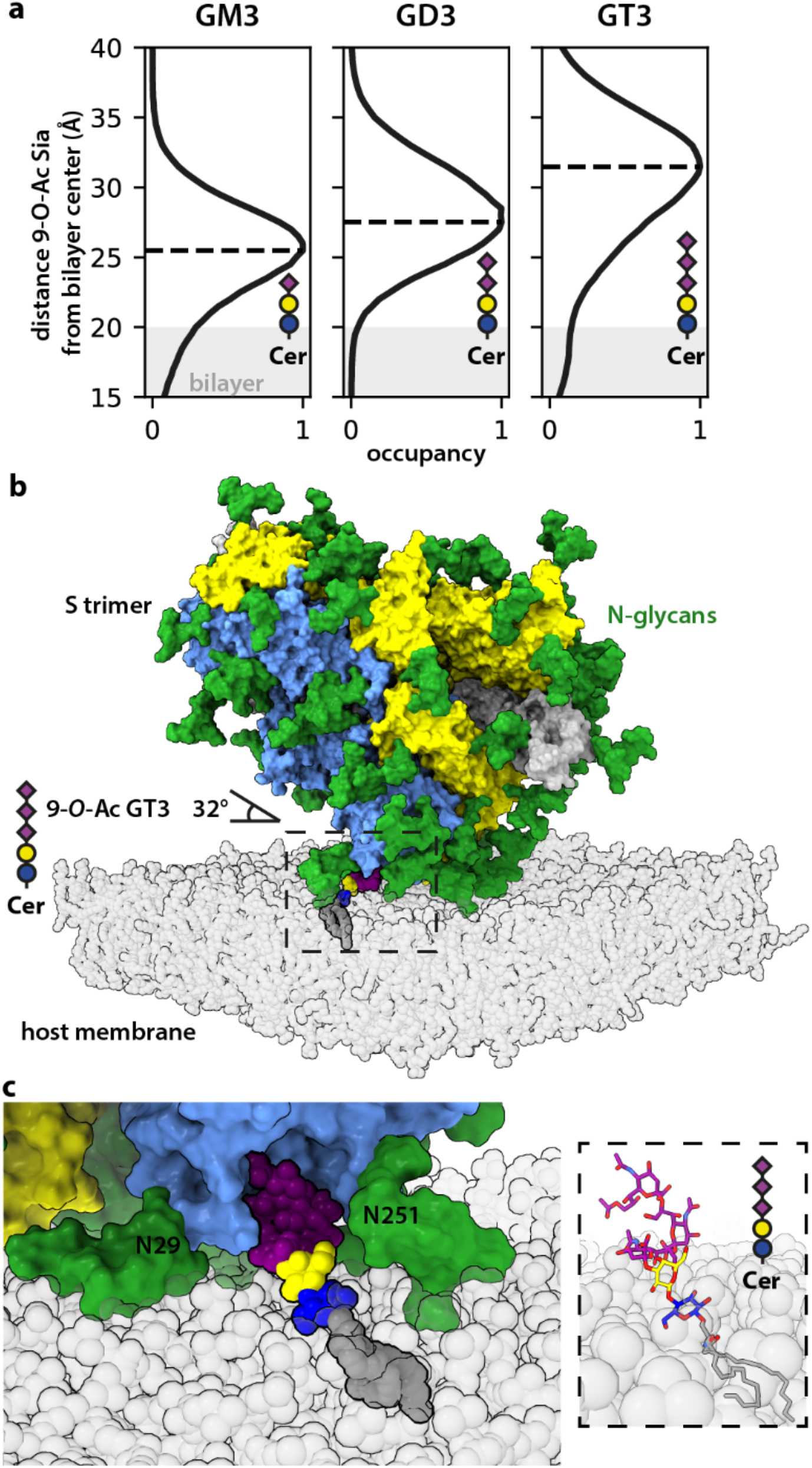
**a.** Lengths by which glycan moieties of gangliosides GM3, GD3, and GT3 extend from a phospholipid bilayer as determined by MD analysis. The outer leaflet (with its boundary marked by the lipid head groups) is indicated in gray. **b**, **c**. MD modelling of HKU1 S binding to membrane-embedded GT3. Binding would require N-glycans at positions 29 and 251 to adopt conformations parallel to the membrane surface. Phospholipids are shown in semi-transparent ball representation.

## DISCUSSION

### Spike glycosylation controls disialoside receptor selectivity in human coronavirus HKU1

Circulation of HKU1 under continuous pressure of the host immune system has selected for a two-lineage evolutionary trajectory. While there is evidence for high-frequency interstrain recombination^2^, HKU1 strains can nevertheless be divided into two distinct serotypes based on diversity in their envelope proteins^3,13^. Given the substantial sequence divergence in these proteins, the serotypical split seemingly occurred early after HKU1 emergence, prompting the question whether differences between serotypes are limited to antigenicity or whether biological differences may also exist. Here, we studied an HKU1 serotype B S protein, focusing on receptor binding properties, the structural basis for S1^A^ ligand specificity and the implications for natural infection. We show that the spike of HKU1-B strain N5, like those of HKU1-A strains Caen1^13^ and N1^15^, preferentially binds to 9-*O*-acetylated disialoside moieties with dissociation constants in the 100 µM range. Despite the evolutionary distance separating A- and B-type spikes and substantial sequence diversity in S1^A^, the extended binding site, recently delineated for A-type strain N1, is conserved. Whereas the terminal 9-*O*-acetylated residue docks into its dedicated Sia2 subsite through canonical interactions previously described^13,21,39^, the penultimate Sia residue is well-accommodated within its own Sia1 subsite through protein-glycan interactions with the e2 element and adjacent residues. The importance of Sia1 for binding and its critical contribution to binding affinity was demonstrated by structure-guided, single-site substitutions resulting in loss or gain of function. Importantly, however, we find that selective receptor-binding is not based on protein-ligand interactions exclusively. The conserved N-glycans at Asn29 and Asn251, decorating the e1 and e2 elements, not only modulate binding affinity (Fig. 3) but also contribute to ligand selectivity. These findings complicate the interpretation of prior observations regarding binding affinity^15^ or ligand specificity^27^, obtained with Man5-glycosylated S produced in N-acetylglucosaminyltransferase I-deficient HEK293 cells. Site-specific glyco-analyses of soluble S trimers of SARS-CoV-2 and HKU1-B N5 expressed in wild-type HEK293 cells found underprocessed oligomannose-type sugars only as minority populations restricted to specific sites. The majority of N-linked glycans were processed (68% and 75%, respectively), with complex-type N-glycans attached to HKU1 residues Asn29 and Asn251^28^. While there are no data available for the processing states of the glycans of virion-associated HKU1 S, the glycosylation profiles of native viral spikes of SARS-CoV, MERS-CoV and SARS-CoV-2 closely resemble those of corresponding recombinant glycoproteins^28,38,40,41^. Given the apparent regulatory role of the HKU1 S Asn29 and Asn251 glycans in receptor-binding affinity and ligand selectivity, the theoretical possibility exists of changes in the host cell glycosylation machinery during the course of the infectious cycle yielding swarms of progeny virus particles that are functionally diverse with respect to receptor usage. The evolutionary conservation of a binding site tailored for disialoside-based receptors implies a selective advantage during replication in the human airways. Preferential binding to these receptors may have arisen as an adaptation to the human respiratory glycome. In support, an AlphaFold 3 model^42^ of the S1^A^ domain of *Rattus argentiventer* CoV - HKU1’s closest known murine relative - predicts the presence of a Sia2 binding site, but a much smaller e2 ridge lacking a six-amino-acid element (S^246^SNTDN^251^), critically involved in protein-Sia1 interactions as shown by us and others^15^ (Fig. S23). Preferred binding to disialoside motifs, common to B- and C-type gangliosides^31,32^, does not exclude low-affinity binding to monosialylated glycans attached to secretory and cell surface proteins such as are abundantly present in the mucus and glycocalyx^30^. Although we did not detect binding of N5 S to α2,3- or α2,6-linked 9-*O*-Ac-Sias and only modest binding of Caen1 S to α2,3-linked 9-*O*-Ac-α2,3-sialyllactose, low-affinity binding may still occur and may be detectable only upon multivalent presentation of S lectin domains to increase avidity such as in the context of the virion^21,27^. MD simulations suggest that the HKU1 S1^A^ domain can accept any 9-*O*-Ac-Sia irrespective of glycosidic linkage type, albeit without supporting contributions from the penultimate residue. During natural infection, hetero-multivalent, dynamic binding of virus particles to such low-affinity monosialoside ligands in combination with HE-mediated receptor destruction may promote virion motility through the mucus and glycocalyx, in an analogous fashion as suggested for influenza virus^43^. Once at the cell surface, virion binding to 9-*O*-acetylated disialoside moieties of gangliosides would result in a more stable, high-affinity attachment. Limited accessibility of such receptors to the receptor-destroying sialate-O-acetylesterase activity of HE, owing to the S-HE size difference^44^, could allow prolonged association with host cells, providing virions an opportunity to find and attach to their secondary entry receptor TMPRSS2.

### Ligand-induced S conformational changes: not an open and shut case

We previously found spikes of HKU1-A strain Caen1 to transition to open S1^B^-up conformations upon ligand binding by S1^A^, thus exposing the TMPRSS2-binding site^13^. These conformational changes appeared subject to formation of the 9-*O*-Ac-Sia binding site, whether by induced fit or conformational selection, and a register shift of the e1 conformational ‘relay’ element^13,15^. Conversely, our current analysis of the HKU1-B N5 S apo structure found the majority of protomers with the ligand binding site already formed and the ‘relay’ element shifted in register, seemingly primed for opening. Nevertheless, we did not detect spontaneous opening and, more remarkably, did not observe transition to the open state for the ligand-bound complex, not even after prolonged incubation at 37 °C. Our observations, for near-native, minimally modified type A and B S ectodomains with only the S1/S2 cleavage site abolished, differ from those of others who reported spontaneous, ligand-independent opening of the receptor-binding domain both for A- and B-type spikes, including N5^15,17,18^. The discrepant findings may be explained by methodological differences. Theoretically, in the case of HKU1-B N5 S our methods may somehow have selected for conformationally-stable subpopulations. We note, however, that in each of the conflicting studies, S ectodomains with stabilizing mutations were used, including consecutive Pro substitutions of S2 residues Asn1067 and Leu1068. The latter S-2P mutations are located in a loop, connecting the S2 central helix (CH) and heptad repeat 1 (HR1) that are at the core of the S fusion machinery. These substitutions aim to restrict the rearrangement of CH and HR1 into an extended α-helix, thereby preventing the S trimers from transitioning from a prefusion to postfusion conformation^22^. Saliently, however, the CH-HR1 helix-turn-helix elements (HTHEs) also form the interface between S2 fusion stalk and S1^B^ domains with inter-domain hydrogen bonding potentially adding to the interaction (Fig. S11). These structural contacts would make an effect of the 2P substitutions on S1^B^ dynamics plausible and might inadvertently facilitate a down-to-up transition of S1^B^. Indeed, 2P substitutions have been shown to affect S1^B^ dynamics in the case of SARS-CoV-2 S^24^. Of note, our findings for the HKU1-B N5 apo structure are consistent with those of Kirchdoerfer et al.^16^. Moreover, a minimally modified version of the OC43 S ectodomain complexed with 9-*O*-Ac-Sia also remained fully closed^39^.

All things considered, our present findings may well reflect a genuine difference between A- and B-type HKU1 spikes. If so, for HKU1-B spikes, prying the lid open may require yet unknown factors beyond primary receptor-binding by S1^A^ or conditions other than ligand-induced local conformational changes alone. Hypothetically, torque acting on virus particles when bound to membrane-embedded sialoglycolipids might provide required mechanical assistance to promote S1^B^ release. From the apparent accessibility of the HKU1-B S1^B^ receptor-binding site in the closed conformation, even the formation of a ternary ganglioside-S-TMPRSS2 complex may be considered as a prelude to S opening and fusion.

### Targeting of ciliated cells through conserved disialoside recognition: a key determinant of host cell tropism?

A previous study, involving ST8SIA1-overexpression in standard cultured cells and HKU1 S pseudoviruses, implicated gangliosides as potential viral receptors^14^. Following up on these observations, we asked whether these glycolipids may also serve as receptors during natural infection. Using human nasal epithelial cell cultures as a model system, ciliated cells were identified as host cells of HKU1, and their infection was strictly dependent on cell surface expression of 9-*O*-acetylated sialoglycans as was also noted elsewhere^5^. We show that both A- and B-type HKU1 S1^A^-Fc proteins, when used as probes, specifically bind to ciliated cells, particularly to 9-*O*-Ac-sialoglycotopes profusely present on cilia. While we did not detect GD3 or 9-*O*-Ac-GD3 in HNEC cultures, we did find evidence for cell surface GT3-like trisialylated glycolipids, again specifically in ciliated cells. Competitive binding of S1^A^-based probes showed that a considerable proportion of these gangliosides are modified by terminal sialate-*O*-acetylation and thus could serve as virus receptors. In further support for potential usage of 9-*O*-Ac-GT3-like gangliosides for viral attachment, the accessible glycan chain of membrane-embedded GT3, but not GD3, was geometrically compatible with S1^A^-mediated attachment as observed for the free-state S-disialoside complex with both Sia2 and Sia1 accommodated.

Our findings extend our understanding of HKU1 biology. The data suggest that preferential binding to 9-O-acetylated disialosides is a common trait. The biological importance of ligand selectivity during natural infection is indicated by the conservation of an extended binding site in A- and B-type spikes. Rather than mere attachment factors, the glycan-based ligands appear to be genuine primary receptors, attachment to which is essential for infection. As was shown for the HKU1 N5 S protein, S1^A^-ligand binding is a prerequisite for S1^B^-TMPRSS2-dependent membrane fusion and entry^10^. The specific and abundant cell surface expression of 9-*O*-Ac-disialosides in ciliated cells suggests that these glycan motifs may be a key, if not the prime determinant of HKU1 host cell tropism.

## METHODS

### Cloning and constructs

Codon-optimized genes of HKU1-A, isolate Caen1 (GenBank ID ADN03339, residues 14-1266) and HKU1-B, isolate N5 (UniProt ID Q0ZME7, residues 14-1261) were synthesized by GenScript. The ectodomain coding sequences were subcloned into a pCG2 expression vector with an exogenous CD5 signal peptide and a C-terminal GCN4 trimerization motif followed by a thrombin cleavage site and a Strep-tag. The furin cleavage site (residues 756-760 in HKU1-A and 752-756 in HKU1-B) was exchanged to GGSGS. S1^A^ domains (residues 14–298) and bovine CoV HE (in a pCD5 vector, UniProt ID P15776, residues: 19-389) were subcloned in-frame with a thrombin cleavage site, a linker and a human Fc tag. Amino acid sequences for all proteins are given in Supplementary Table S2. Subcloning was performed using NEBuilder HiFi DNA assembly (NEB). Primers were designed using SnapGene (v7) with target melting temperatures of the 15–25-base overlaps of 50 °C. Linear insert and vector fragments were amplified using Q5 (NEB) and Primestar GXL polymerases (Takara Bio), respectively, according to the manufacturers’ instructions. Success of PCR reactions was checked by agarose gel (1% w/v) electrophoresis. PCR products were incubated with DpnI (NEB) for 1 h at 37 °C followed by clean-up using the NucleoSpin PCR Clean-up kit (Macherey-Nagel). Fragments were ligated at a 1:2 molar ratio (vector: insert) with 100 ng vector DNA using NEBuilder HiFi DNA Assembly Master Mix (NEB) according to the manufacturer’s instructions for 1 h at 50 °C. Bacterial transformation was done with 2 µl of ligation mixtures in chemically competent XL10 cells which were spread on LB agar plates containing 100 µg/ml ampicillin. The presence of the respective inserts was established by colony PCR and agarose gel electrophoresis and confirmed by DNA sequencing. For the generation of point mutants, ‘back-to-back’ primers (designed by the NEBaseChanger tool) and Primestar GXL polymerase were used for synthesis of linear DNA. 1 µl of PCR product was treated with KLD mix (0.2 µl T4 ligase, 0.4 µl T4 kinase, 0.4 µl DpnI, 1 µl 10× T4 ligase buffer, 0.5 µl PEG4000, 6.5 µl H_2_O) for 1 h at room temperature before bacterial transformation.

### Protein expression and purification

HEK293T and HEK293S GnTI-cells were cultured in DMEM high glucose with L-glutamine supplemented with 10% v/v FCS and penicillin/streptomycin (all Capricorn Scientific) at 37 °C and 5% CO_2_. For protein expression of trimeric S ectodomain and S1^A^-Fc constructs (see Supplementary Table S2), 2 × 10⁷ cells were seeded in T225 flasks. After 24 h, cells in each flask were transfected with a mixture of 200 μg PEI-25000 (transfection grade, 1 mg/ml stock solutions adjusted to pH 7, Polysciences Inc.) and 40 μg of the expression plasmid dissolved in 2 ml serum-free DMEM for 6-16 h. The culture medium was then exchanged to serum-free medium (per liter of 293 SFM II medium [Thermo Fisher Scientific]: 10 ml GlutaMAX [Gibco], 15 ml DMSO, 2 g glucose, 3.7 g NaHCO_3_, 3 g peptone primatone [Sigma]). Culture supernatants were harvested after 5-7 d and clarified by centrifugation at 20,000 g for 1 h at 4 °C and filtration with a 0.2 μm bottle-top filter (Corning). A solution of 1 M Tris, pH 8.8, was used to adjust the supernatant to pH 7. All proteins were purified on an AKTA Go system (Cytiva) by monitoring UV absorption at 280 nm. Supernatants containing Strep-tagged trimeric S ectodomains were applied to 1 ml StrepTactin columns (Cytiva) at a flow rate of 1 ml/min and washed with 0.1 M Tris, 10 mM NaCl, 1 mM EDTA (pH 8). Proteins were eluted using 50 mM D-biotin (Carl Roth) in washing buffer. The eluate was applied to a Superose 6 Increase size exclusion column (Cytiva) in 20 mM sodium phosphate buffer, 100 mM NaCl (pH 7.3). Human Fc-tagged S1^A^ domains were affinity-purified in a similar manner on 1 ml Protein A columns (Cytiva). Proteins were eluted in 300 μl fractions using 0.1 M citric acid, pH 3, into 96-well plates containing 100 μl 1 M Tris, pH 8.8. Size exclusion chromatography was performed using a Superdex 200 Increase column (Cytiva). All proteins were concentrated using 10 kDa MWCO centrifuge filters (Merck), flash-frozen in liquid nitrogen and stored at −80 °C. All protein concentrations refer to the respective monomeric proteins.

Where indicated, Fc-fusion proteins were cleaved using 50 U thrombin per mg of protein during dialysis against 10 mM Tris, 50 mM NaCl (pH 8) for 16 h at room temperature. Remaining uncleaved proteins and cleaved Fc fragments were removed by protein A chromatography as described above. The unbound fractions containing monomeric S1^A^ domains were subjected to size exclusion chromatography using a Superdex 75 Increase column (Cytiva).

### Human airway epithelial cell culture

Primary human nasal epithelial cells (HNECs) from consenting healthy donors (identifiers 0266 and 0263) were obtained from the University Medical Center Utrecht Biobank, Utrecht, the Netherlands, with permission from the UMC Utrecht Biobank Research Ethics Committee (TCBio 21-265). Cells were collected by nasal brushing, propagated into working cell-bank stocks and maintained as described previously^29^. In brief, primary cells were propagated in collagen IV-coated flasks (Sigma-Aldrich) using basal cell (BC) expansion medium, refreshing three times per week until there were sufficient numbers of cells to seed onto collagen IV-coated 24-well polyethylene terephthalate, 0.4 µm pore Transwell inserts (Corning). Cells were cultured in submerged conditions with BC expansion medium in both the basolateral and apical compartments until confluent. Basolateral and apical media were then replaced with ALI-differentiation medium with additional TGF-β inhibitor A83-01 (500 nM final concentration). After 2 days, basolateral medium was refreshed with ALI-differentiation medium with additional A83-01, apical medium was removed and cells were washed once in Dulbecco’s phosphate buffered saline (PBS) without calcium and magnesium (10 min, 37 °C, 5% CO_2_). After a further 2 days, ALI-differentiation medium without additional A83-01 was used in the basolateral compartment. Cultures were maintained for 21 days in air-exposed conditions, replacing the basolateral medium twice per week and conducting an apical wash in PBS without calcium and magnesium (10 min, 37 °C, 5% CO_2_) once per week to remove any overlying mucus. Successful differentiation was determined by the presence of beating cilia under a light microscope.

### Immunofluorescence assays and virus infections

The cell culture medium was removed from the basolateral compartment of HNEC Transwell inserts and cells were fixed by addition of 3.7% formaldehyde to both sides and incubation for 30 min, followed by three washing steps with PBS. Controls were treated with bovine CoV HE (50 μg/ml) or neuraminidase from *Arthrobacter ureafaciens* (100 mU/ml, Merck) for 2 h at 37 °C and washed twice with PBS. Methanol treatment was performed with ice-cold methanol for 15 min and lipid extraction by consecutive submersion of Transwell membranes in the following solutions for 5 min each: H_2_O, 70%, 96%, and 100% EtOH, xylene, 100%, 96%, and 70% EtOH, H_2_O. All cells were blocked with 3% BSA, 0.05% Tween-20 in PBS for 1 h. All staining steps were performed with reagents diluted in blocking buffer for 1 h at room temperature and three subsequent washing steps. S1^A^-Fc proteins and anti-ganglioside antibodies were used at concentrations of 50 μg/ml and 2.5 µg/ml, respectively, unless stated otherwise (anti-A2B5: ab53521 Abcam, UM4D4: sc-32269 Santa Cruz Biotechnology, R24: MABC1112 Merck). Biotinylated *Sambucus nigra* lectin was used at a dilution of 1:100 (B-1305-2 Vector Labs). Non-permeabilized cells used for surface staining were stained with the following secondary antibodies: Donkey Anti-Human IgG AF488 (1.5 µg/ml, 709-545-149 Jackson ImmunoResearch), Goat anti-Mouse IgM AF594 (1:400, A-21044, Thermo Fisher Scientific), Goat anti-Mouse IgG AF594 (1:400, A-11005 Thermo Fisher Scientific), and streptavidin AF568 (1:500, S11226, Thermo Fisher Scientific). Co-staining of surface-stained cells for β-tubulin was preceded by a second fixation step with 3.7% formaldehyde for 30 min, two washing steps with PBS, and permeabilization with 0.5% Triton X-100 in blocking buffer. Staining was performed using a rabbit anti-β-tubulin antibody (1:200, ab6046 Abcam), and Donkey anti-Rabbit IgG AF647 (1:200, A-31573 Thermo Fisher Scientific). Secondary staining solutions contained 0.1 µg/ml DAPI (Sigma-Aldrich) and, where indicated, phalloidin AF568 (1:50, A-12380 Thermo Fisher Scientific). Transwell membranes were mounted on microscopy slides using ProLong Diamond Antifade Mountant (Thermo Fisher Scientific).

HKU1-A virus, isolate Caen1, was kindly provided by Krzysztof Pyrc (Jagiellonian University Krakow). Before infections, mucus was removed from differentiated HNECs by washing three times with PBS. For the initial passage, infections were performed at 33 °C with 8 µl of virus stock (5.37 × 10⁶ RNA copies/ml^19^) in 100 µl PBS per Transwell insert. Mock-infected cells were treated with PBS only. The inoculum was removed after 1 h followed by three washing steps. Cells were maintained at 33 °C for 96 h. A 1:4 dilution of an apical wash with 100 µl PBS collected at 96 h p.i. was used for a second round of infections. Control cells were treated with bovine CoV HE (390 µg/ml) for 1 h at 37 °C prior to infection. Cells were fixed at 24 h p.i. and kept in PBS at 4 °C before staining. Fixed cells were permeabilized using 0.5% Triton X-100 in blocking buffer for 1 h. Staining reactions were performed as described above with primary staining solutions containing rabbit anti-MHV polyclonal serum K135 (1:300^45^) and mouse anti-β-tubulin (1:200, MU178-UC Biogenex) or mouse anti-dsRNA (1:500, J2 Scicons) with rabbit anti-β-tubulin (1:200, ab6046 Abcam). Secondary staining solutions contained Donkey anti-Rabbit IgG AF647, Goat anti-Mouse IgG AF488 (1:400, A-11001 Thermo Fisher Scientific), phalloidin AF568, and DAPI.

All samples were imaged on a Nikon A1R confocal microscope using NIS-Elements (v4.5). Z-stack sizes, pinhole diameters, objective lenses, filters, and image sizes are specified in Supplementary Table S3. The scan speed was set to 0.5 and two-fold line averaging was applied. Images were processed using Fiji^46^ with ImageJ v1.54 and colored according to color-blind-accessible palettes. Surface-stained samples are shown as maximum intensity projections unless stated otherwise. 3D views were generated by ‘tricubic sharp’ interpolation. The number of infected cells was assessed from 3 × 3 fields of view stitched with 15% overlap using a thresholding value of 400 for 16-bit images and ImageJ’s ‘count object’ function with a minimum area of 25 µm². Standard deviations across three overview images are given as error bars.

### Biolayer interferometry

BLI experiments were performed on an Octet Red384 (ForteBio) at 25 °C. Biotinylated sialoglycan ligands^14^ were immobilized on SA biosensors (Sartorius) at a concentration of 2 μM in 20 mM sodium phosphate buffer, 100 mM NaCl, pH 7.3 (PB) for 180 s unless stated otherwise. Sensors were transferred into blocking buffer (BB, 1 mg/ml BSA, 0.05% v/v Tween-20 in PB) for 180 s. S1^A^-Fc proteins and trimeric S ectodomain samples in PB were adjusted to BB conditions by addition of a 20× BSA/Tween-20 stock. Analyte binding was observed for 360 s and 2000 s for S1^A^-Fc proteins and S ectodomains, respectively. Concentrations of S1^A^-Fc constructs were 2.5 µM unless stated otherwise. Response levels were evaluated from two measurements with pristine sensors for S1^A^-Fc proteins and S ectodomains at the end of the association phase and the respective standard deviations are given as error bars. Monomeric S1^A^ domains in a concentration range of 6.25–95 µM were used for determination of binding kinetics and dissociation constants. Association and dissociation in BB were observed for 120 s each for three cycles. Data were aligned with the ForteBio data analysis software (v9) and then averaged and used for curve fitting to a 1:1 binding model (equations 1–2) using in-house Python scripts and the lmfit library^47^. Kinetic rate constants were used as global fit parameters.

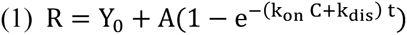

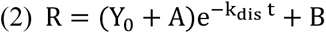

Y_0_ and B denote the association- and dissociation-phase baseline responses, respectively; A denotes the response amplitude; k_on_ and k_dis_ denote the association and dissociation rate constants, respectively; and C denotes the protein concentration. Steady-state responses (Y_0_+A) derived from the S1^A^ domain concentration series were used for validation of obtained K_D_ values by curve fitting against equation 3. The maximum binding response R_max_ was used as a global fit parameter.

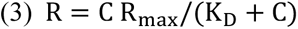

### Cryo-EM sample preparation

HKU1-B S ectodomain samples were prepared in 20 mM sodium phosphate buffer, 100 mM NaCl (pH 7.3). The (monomeric) protein concentration was 5 μM for the unbound protein and 4.7 μM for the disialoside complex. 9-O-Ac-Neu5Ac-α2,8-Neu5Ac-Lc-biotin^14^ was titrated from a 5 mM stock solution for a final concentration of 300 μM. Samples were incubated for 10 min at room temperature or for 1 h at 37 °C and applied to QuantiFoil R1.2/1.3 grids that were glow-discharged for 20 s at 20 mA. Blotting and plunge freezing were performed in a Vitrobot Mark IV (Thermo Scientific) for 5 s at 20 °C and 95% relative humidity with a blot force of 0.

### Cryo-EM data collection

Cryo-EM data for the apo and holo datasets were acquired on a Krios G4 transmission electron microscope (Thermo Fisher Scientific) operated at 300 kV and equipped with a Selectris X energy filter (Thermo Fisher Scientific) and a Falcon 4i direct electron detector (Thermo Fisher Scientific). Movies were recorded in EER mode using EPU software (Thermo Fisher Scientific) at a nominal magnification of 165,000×, corresponding to a calibrated pixel size of 0.73 Å. The energy-filter slit width was set to 10 eV. Data were acquired with a stage tilt of 32°, an exposure time of 3.46 s and a total electron exposure of approximately 50 e⁻/Å². In total, 2,445 apo and 5,402 holo movies were collected. The 37 °C-incubated dataset was acquired at Utrecht University on a Talos Arctica transmission electron microscope (Thermo Fisher Scientific) operated at 200 kV and equipped with a K2 Summit direct electron detector (Gatan) and a post-column GIF Quantum energy filter (Gatan). Movies were recorded in counting mode using EPU software (Thermo Fisher Scientific) at a nominal magnification of 130,000×, corresponding to a calibrated pixel size of 1.04 Å. The energy-filter slit width was set to 20 eV. Data were acquired with a stage tilt of 32°, an exposure time of 10.0 s and a total electron exposure of 50 e⁻/Å². In total, 510 movies were collected.

### Cryo-EM data processing

Data-processing schemes and parameters for the apo and holo HKU1-B datasets are shown in Figs. S3 and S5, respectively. cryoSPARC v4.4 was used for all data processing^48^. Patch motion correction, patch CTF estimation and initial particle picking using the template-free blob picker were performed in cryoSPARC Live. An initial round of 2D classification was used to generate templates for a second round of particle picking, followed by particle curation, extraction, 2D classification and ab-initio reconstruction. Closed spike classes identified by ab-initio reconstruction were selected and re-extracted in 500-pixel boxes and Fourier-cropped to 340 pixels, corresponding to a sampling of approximately 1.074 Å per pixel. The selected apo and holo particle stacks contained 61,254 and 196,929 particles, respectively, and were subjected to C3 non-uniform refinement^49^. Reference-based motion correction was subsequently performed with re-extraction at the original sampling of 0.73 Å per pixel in 500-pixel boxes. Final C3 non-uniform refinements contained 61,233 apo and 196,925 holo particles and yielded global maps at resolutions of 2.57 Å and 2.40 Å, respectively, according to the gold-standard FSC 0.143 criterion^50^.

For focused refinement of the ligand-binding region, the holo particle stack was C3 symmetry-expanded to 590,775 particles. A mask encompassing one S1^A^ domain was generated in UCSF Chimera^51^ and used for local refinement, yielding a map at 2.74 Å resolution according to the gold-standard FSC 0.143 criterion. Local-resolution estimation and filtering were performed for the apo and holo global reconstructions and the focused S1^A^ reconstruction. cryoSPARC 3D variability analysis^23^ was used to visualize conformational heterogeneity and domain motions within S1^A^. The local-resolution-filtered global maps and focused S1^A^ map were used for model building.

For the spike–disialoside complex incubated for 1 h at 37 °C, following template-based particle picking and 2D classification, 59,146 selected particles were subjected to ab-initio reconstruction. Particles were subsequently re-extracted in 500-pixel boxes and Fourier-cropped to 300 pixels, subjected to heterogeneous refinement and further 2D classification, and combined for a final C3 non-uniform refinement containing 58,585 particles. The resulting reconstruction had a global resolution of 5.57 Å according to the gold-standard FSC 0.143 criterion (Fig. S9).

### Model building

A structure of HKU1-B S (PDB ID 5I08) was used as the starting model for building the apo and disialoside-bound structures into their respective local-resolution-filtered global maps. The focused S1^A^ map was additionally used to model the ligand-binding region of the holo structure. Initial phases of model building were performed iteratively using Coot v.0.9.8.7^52^ and real-space refinement in Phenix v.1.20.1^53^ as distributed by SBGrid. N-glycans were modelled using Coot’s carbohydrate module where supported by experimental density. Final model building steps to resolve geometrical conflicts were performed using Isolde^54^ as implemented in UCSF ChimeraX v1.6^55^ followed by a final round of real-space refinement in Phenix. MolProbity^56^ and Privateer^57^ were used for model validation of the protein and N-glycans, respectively. For modelling of the ligand, two unlinked sialic acid residues were first placed inside the respective densities and fitted using the Isolde environment. Atoms of the α2,8-linkage were modelled according to previous findings on allowed regions of the four relevant dihedral angles^13^. Backbone RMSD values were calculated using the jFATCAT (rigid) algorithm as implemented in the RCSB PDB alignment tool^58^.

### Molecular dynamics simulations

Starting structures of the molecular systems were built based on the cryo-EM structure of the HKU1-B S protein (this work) using the graphical interface of YASARA^59,60^. The N-glycans were attached to the protein based on data from quantitative site-specific N-linked analysis of HKU1 S^28^. The full carbohydrate ligands were positioned manually into the binding site located in the NTD guided by the α2,8-linked 9-*O*-acetyldisialoside motif present in the *holo* cryo-EM structure. Starting structures of the 9-*O*-acetylated gangliosides (GM3, GD3, GT3) embedded into a POPC model membrane were also built using the graphical interface of YASARA. The POPC membrane size was 50 Å × 50 Å and the simulation time was 1 µs for GM3 and GD3 and 5 µs for GT3. The structure of the membrane-bound HKU1 trimer was built in two steps. First, a complex of HKU1 trimer with 9-O-Ac-GT3-Cer was modeled and the ceramide tail was adjusted to point away from the protein surface. In a second step the complex was embedded into a POPC membrane (208 Å × 208 Å).

In general, the systems were solvated in 0.9% NaCl solution (0.15 M) and simulations were performed at 310 K and 1 bar under periodic boundary conditions using the AMBER14 force field^61,62,63,64^. The simulation box of the HKU1 trimer bound to GT3 in a POPC membrane contained 962,920 atoms and could be simulated with a performance of 3 ns/day using YASARA with GPU acceleration in ‘fast mode’ (4 fs time step)^60^ on ‘standard computing boxes’ equipped, for example, with one 12-core i9 CPU and NVIDIA GeForce GTX 1080 Ti. Conformational Analysis Tools (CAT)^65^ was used for analysis of trajectory data, general data processing and generation of scientific plots.

### Analysis and visualization

Protein–protein and protein–ligand interactions were analyzed using PDBePISA^66^ and LigPlot+^67^. Figures were generated using UCSF ChimeraX^68^. Structural biology applications used in this project were compiled and configured by SBGrid^69,70^.

## Data availability

Atomic coordinates and cryo-EM maps will be deposited in the Protein Data Bank and Electron Microscopy Data Bank. Correspondence regarding structural data should be addressed to Daniel L. Hurdiss, and requests regarding virology and wet-lab data should be directed to Raoul J. de Groot.

## Acknowledgements

We thank Krzysztof Pyrc for providing the HKU1-A Caen1 isolate, Richard Wubbolts of the Center for Cellular Imaging (CCI) for valuable advice on imaging, Lia van der Hoek (Amsterdam UMC) for insightful discussions, and Jolanda de Groot-Mijnes for critically reading the manuscript. Confocal images were acquired at the Center for Cellular Imaging, Faculty of Veterinary Medicine, Utrecht University. R.C. received funding from the Deutsche Forschungsgemeinschaft (DFG, German Research Foundation; grant 494746248). D.L.H. is supported by the EMBO Young Investigator Programme (YIP-6255) and acknowledges funding from the Beijerinck Premium of the M.W. Beijerinck Virology Fund, Royal Netherlands Academy of Arts and Sciences (KNAW). This project has received funding from the European Union’s Horizon 2020 research and innovation programme under the Marie Skłodowska-Curie Actions grant agreement no. 812673 (OrganoVIR). We thank members of the Utrecht Virology Lab for valuable feedback during preparation of the manuscript and the staff of the Utrecht University Electron Microscopy Centre for technical assistance.

## Author contributions

R.C., D.L.H. and R.J.d.G. conceived the project and developed the overall experimental strategy, with contributions from L.E.W. and R.J.G.H. to experimental design. R.C. designed and constructed the expression plasmids, performed protein expression and purification, and carried out the biolayer interferometry experiments. R.C. and L.E.W. designed and performed the infection experiments and fluorescence-microscopy analyses in human epithelial cultures. I.D. performed cryo-EM sample preparation and data collection. O.J.D.-A. assisted with cryo-EM data collection for the 37 °C Arctica dataset. R.C. and D.L.H. processed the cryo-EM data and built and refined the atomic models. M.F. performed the molecular-dynamics simulations. R.C., M.F., D.L.H. and R.J.d.G. analysed, visualized and curated the data. R.C., D.L.H. and R.J.d.G. were responsible for project administration. R.C., R.J.G.H., B.-J.B., F.J.M.v.K., D.L.H. and C.A.M.d.H. acquired funding. I.D., F.J.M.v.K., B.-J.B., J.B., C.A.M.d.H. and M.F. provided equipment, reagents and other resources. R.C., D.L.H. and R.J.d.G. wrote the original draft of the manuscript. All authors reviewed and edited subsequent versions. D.L.H. and R.J.d.G. supervised the project.

