## Supplementary information for "Spike and host glycan determinants of HKU1 airway tropism"

This PDF file includes:

**Figures S1 to S23**

**Tables S1 to S3**

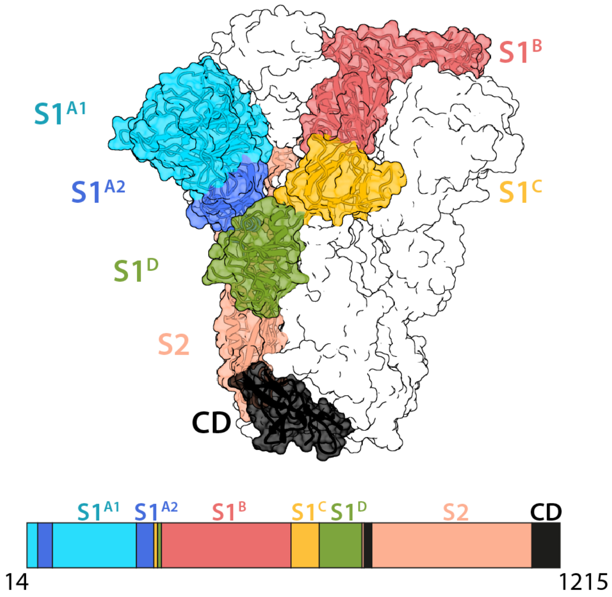


**Fig. S1: S protein domain organization.**

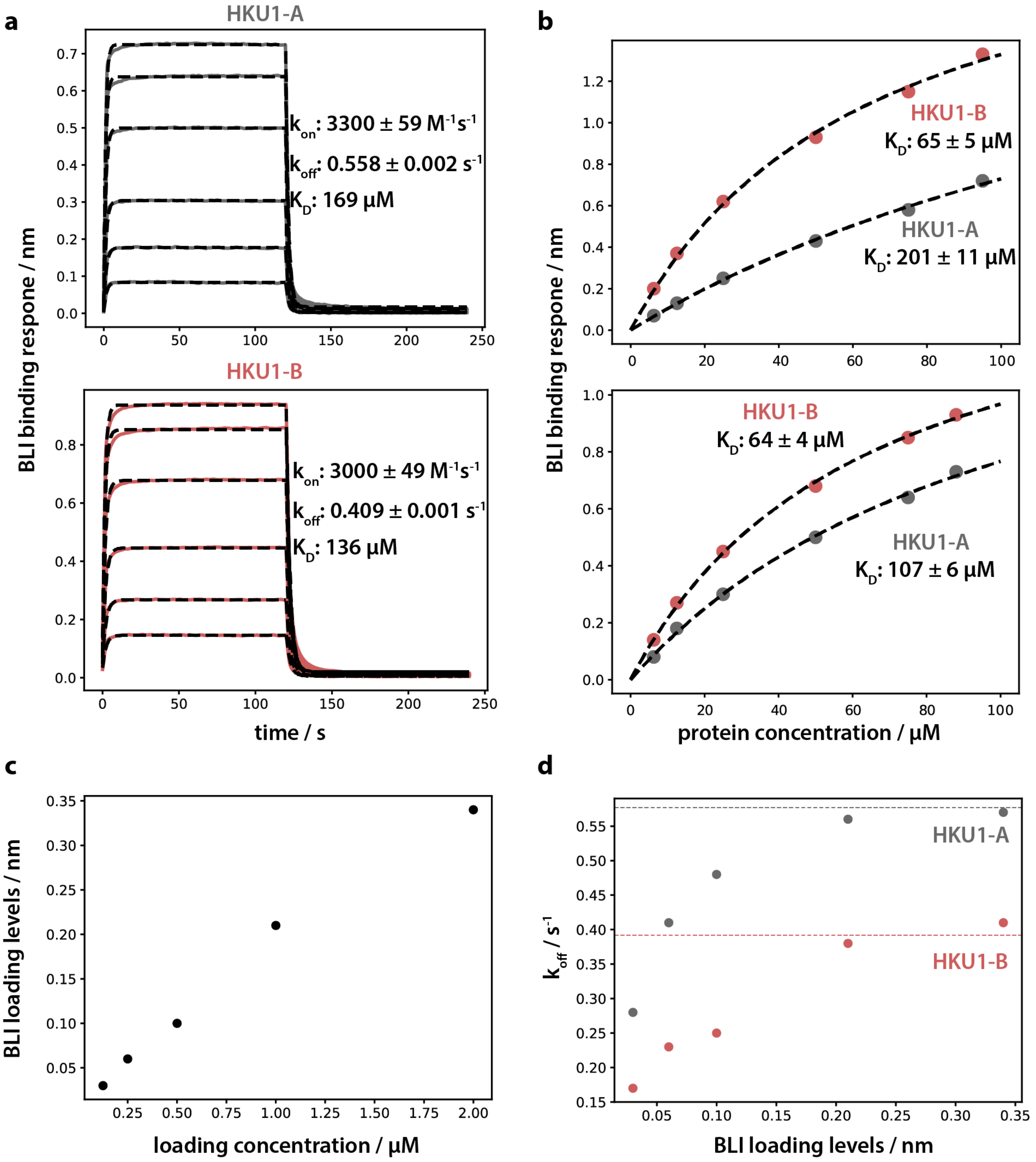


**Fig. S2: Curve fitting of BLI binding data of HKU1-A and -B S1A monomers to 9-O-Ac α2,8-linked disialoside. a.** Global fitting of kinetic binding data to a kinetic 1:1 binding model yielded association and dissociation rate constants. Data shown here are additional replicates of experiments shown in Fig. 1c. **b.** Derived dissociation constants are supported by independent fitting of concentration-dependent steady state binding responses. Curves do not reach saturation levels due to poor protein solubility at higher protein concentrations. **c, d.** Rebinding effects do not decrease HKU1 $S1^A$ monomer dissociation rates. 9-O-Ac α2,8-linked disialoside was immobilized on BLI sensors for 10 s at different concentrations resulting in different loading densities (**c**) below saturation levels. **d.** These sensors (with different amounts of immobilized 9-O-Ac α2,8-linked disialoside) were tested for association and dissociation of HKU1-A and -B $S1^A$ monomers (20 and 8.3 µM, respectively) for 120 s each. Dissociation rates were determined by local curve fitting using ForteBio Data Analysis software v9. Increasing loading levels did not result in a decrease of dissociation rate constants, indicating that rebinding to the surface is not limiting the determination of kinetic binding parameters. Dissociation rate constants determined from global curve fitting are indicated as dashed lines.

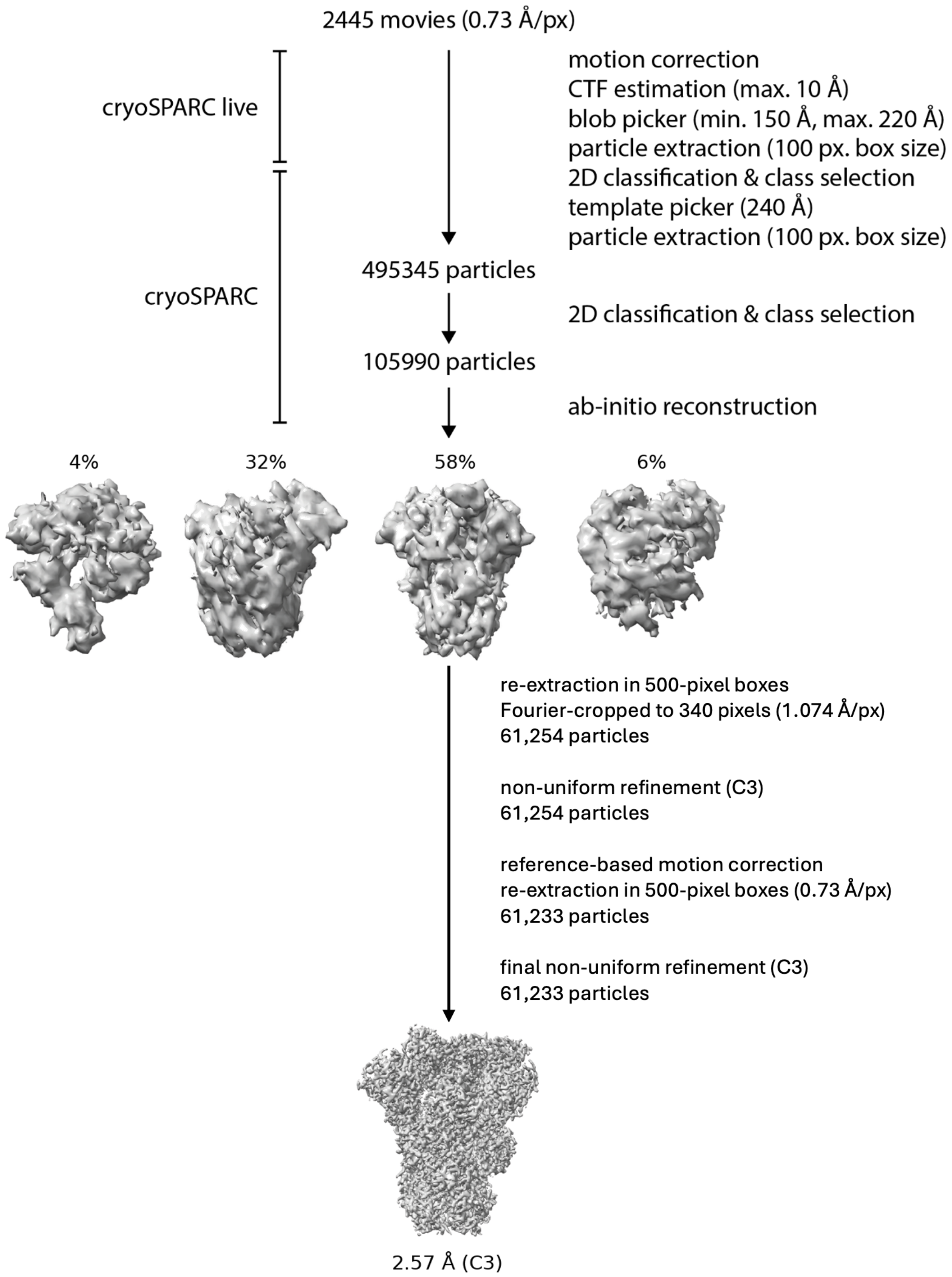


**Fig. S3: Cryo-EM processing pipeline for the unbound '*apo*' HKU1-B S protein data set[1].**

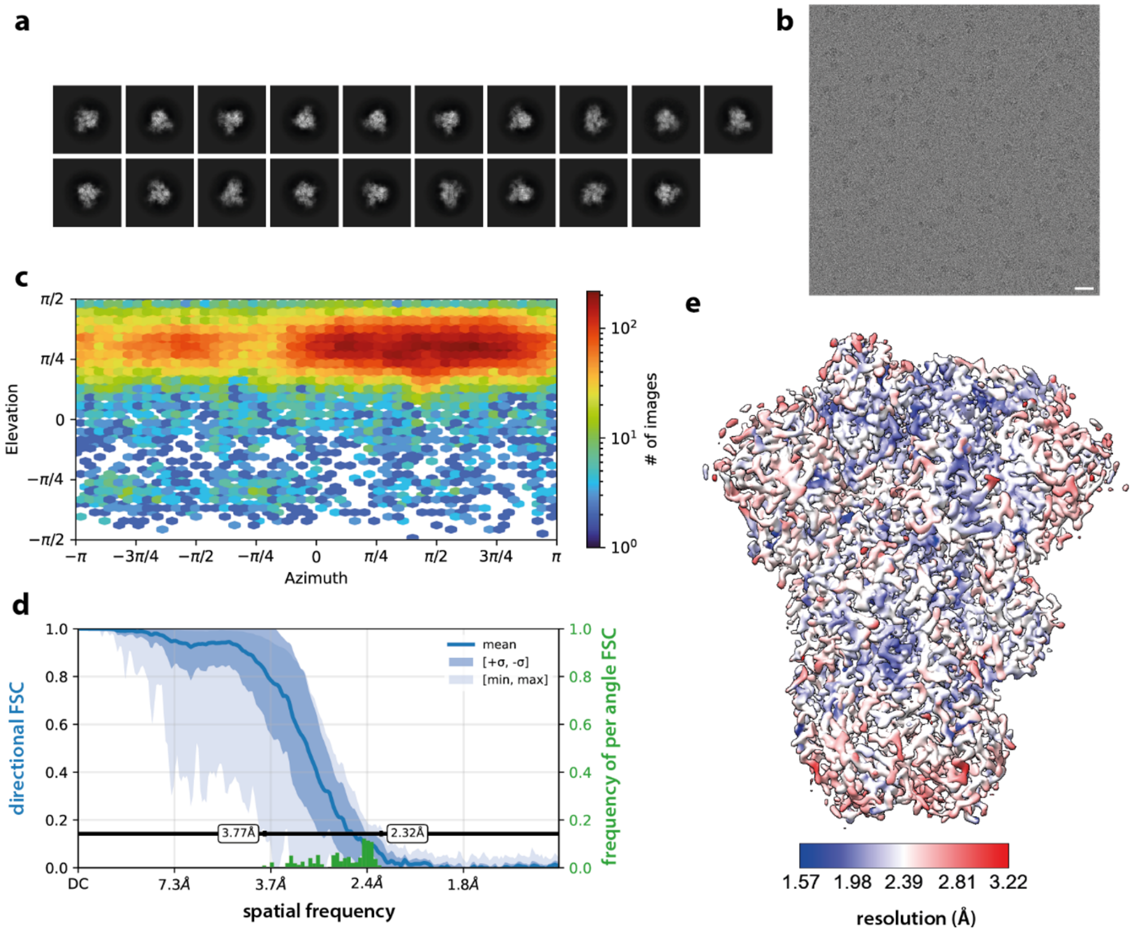


**Fig. S4: Cryo-EM data processing details of unbound HKU1-B S protein. a.** 2D class averages selected for ab-initio reconstruction. **b.** Representative, motion-corrected micrograph of S proteins collected at 32° tilt. The scale bar denotes 200 Å. **c.** Orientation distribution plot for the HKU1-B S map after non-uniform refinement[1]. **d.** 3D FSC analysis[2] showing FSC variation as a function of viewing direction as implemented in cryoSPARC. The histogram over the 0.143 crossings of the directional FSC curves is depicted in green. **e.** Local resolution plotted on the globally refined map of unbound HKU1-B S protein.

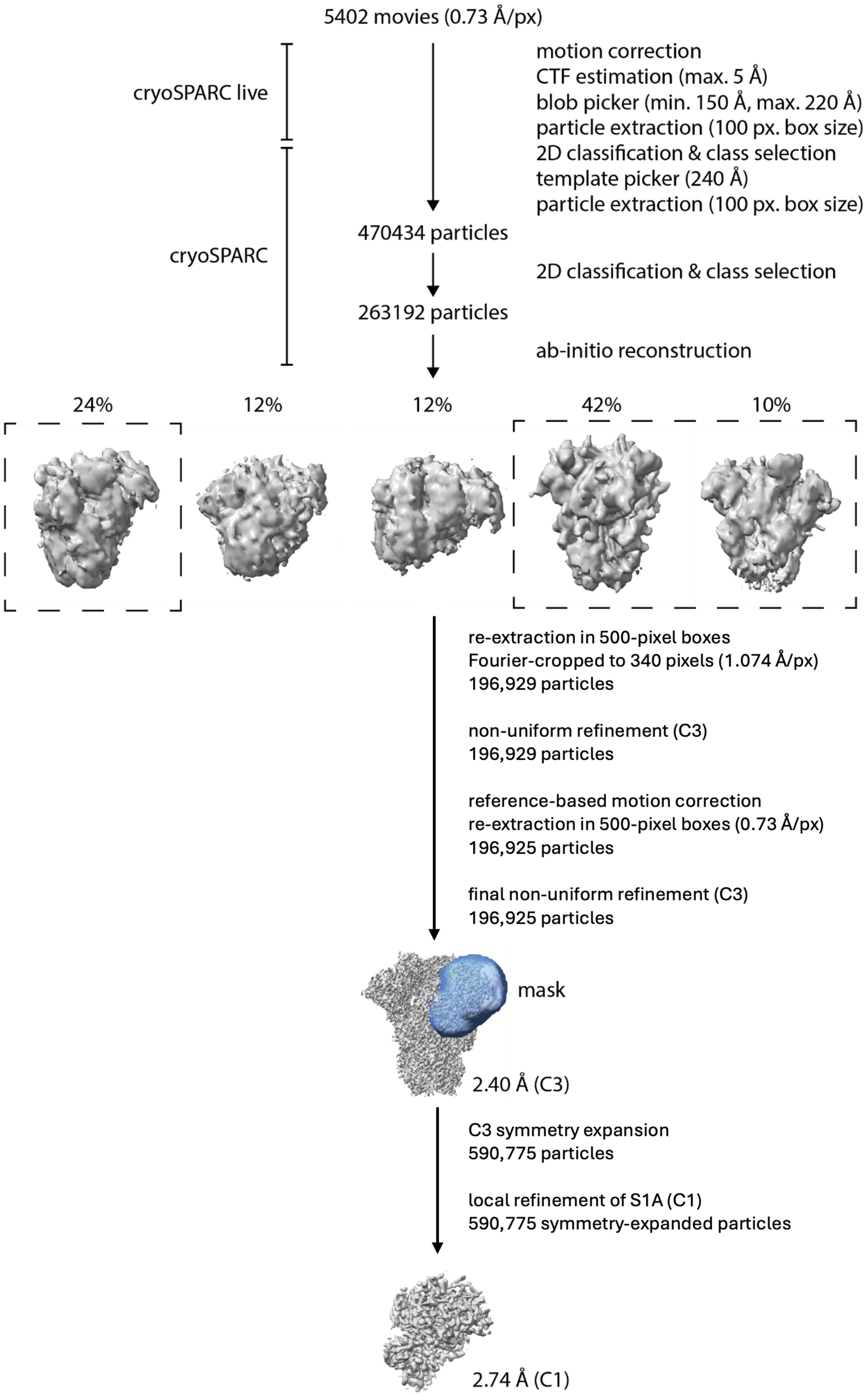


**Fig. S5: Cryo-EM processing pipeline for the HKU1-B S protein in complex with a 9-*O*-Ac disialoside.**[1]

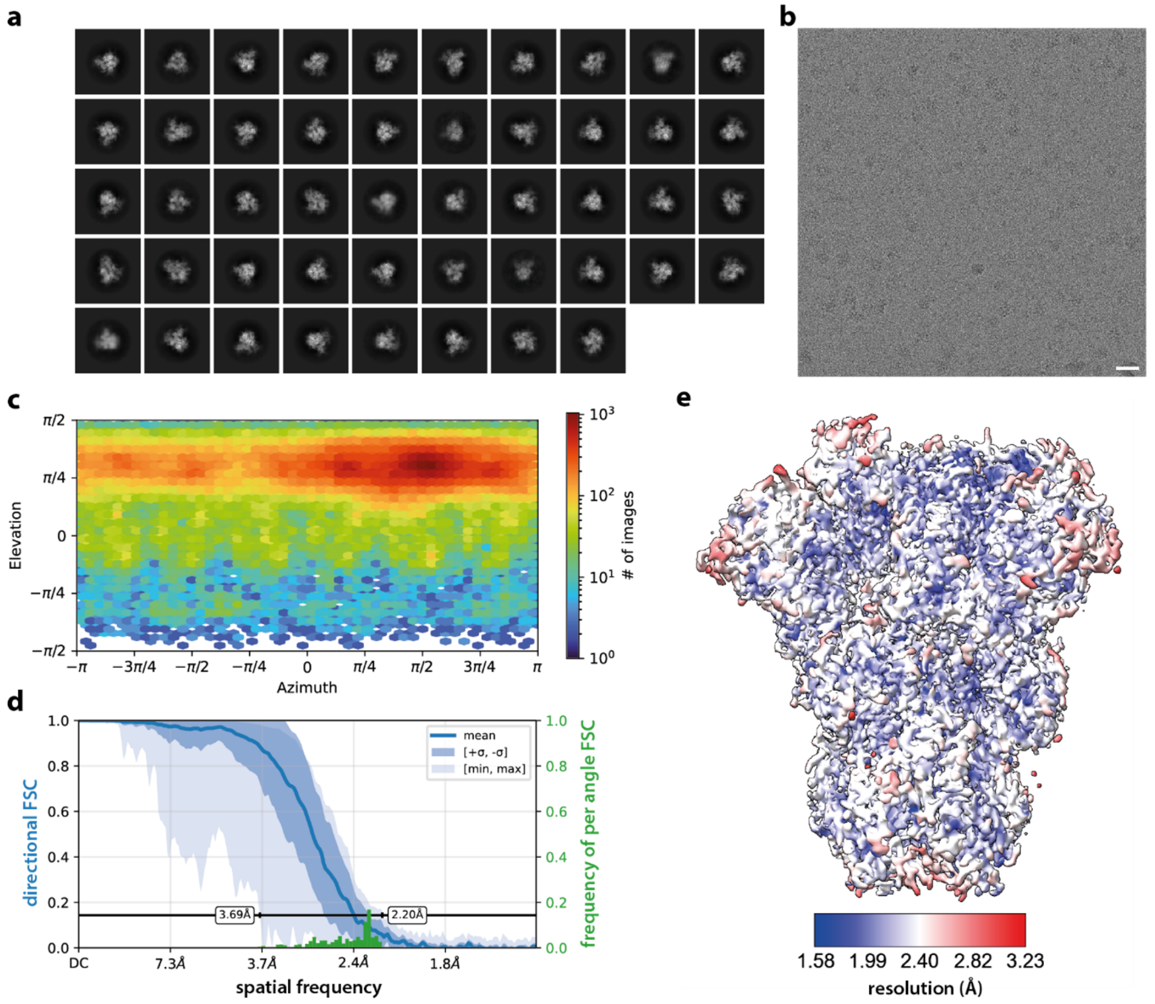


**Fig. S6: Cryo-EM data processing details of HKU1-B S protein in complex with a 9-*O*-Ac disialoside. a.** 2D class averages selected for ab-initio reconstruction. **b.** Representative, motion-corrected micrograph of S proteins collected at 32° tilt. The scale bar denotes 200 Å. **c.** Orientation distribution plot for the HKU1-B S map after non-uniform refinement[1]. **d.** 3D FSC analysis[2] showing FSC variation as a function of viewing direction as implemented in cryoSPARC. The histogram over the 0.143 crossings of the directional FSC curves is depicted in green. **e.** Local resolution plotted on the globally refined map of complexed HKU1-B S protein.

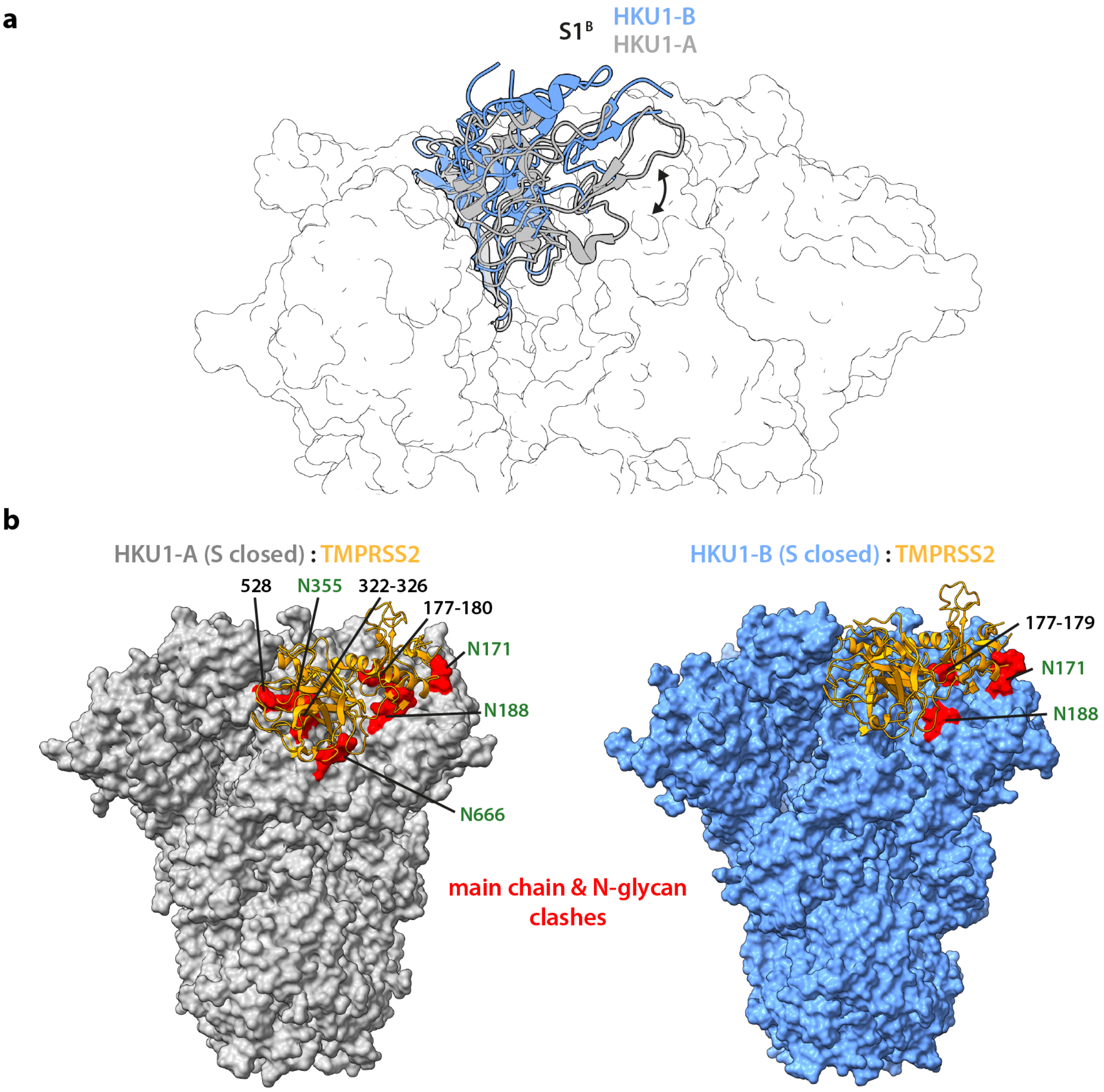


**Fig. S7: Differences in S1$^{B}$ orientation between HKU1-A and -B S proteins. a.** Closed S protein structures of HKU1 serotypes A and B (PDB 8OPM[3] (grey) and the structure determined in this study (blue), respectively) show a different orientation of their respective S1$^{B}$ domains (shown as cartoons), harboring the binding site for TMPRSS2. **b.** Superimposing a recently determined structure of a complex between S1$^{B}$ and TMPRSS2 (orange ribbons, PDB ID 8VGT[4]) with the closed ectodomain S proteins of HKU1-A and -B shows significantly fewer clashes for HKU1-B (clash areas are marked in red). In HKU1-A S, four *N*-glycans and three protein loops interfere with TMPRSS2 binding in the closed state. Many of these clashes are avoided in HKU1-B S due to the more exposed conformation of S1$^{B}$. Clashes were evaluated between main chain atoms or the first carbohydrate residues of *N*-glycans (defined as van der Waals overlap larger than 0.6 Å as implemented in ChimeraX v1.6).

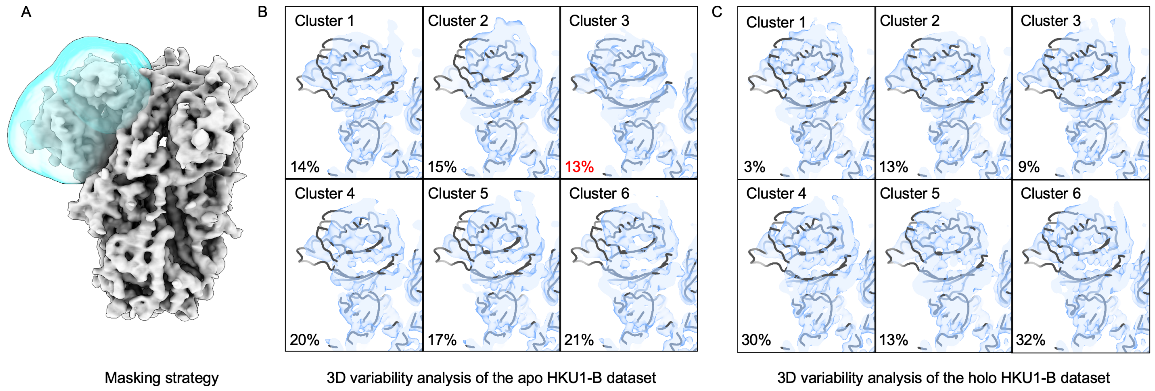


**Fig. S8: Focused three-dimensional variability analysis[5] of apo and ligand-bound HKU1-B spike.** A. Masking strategy used for focused analysis. A soft mask encompassing one $S1^A$ domain and the adjacent $S1^B$ domain is shown in cyan, with the remaining spike density shown in grey. B, C. Six clusters obtained by 3D variability analysis of the apo (B) and 9-O-acetylated-disialoside-bound, holo (C) HKU1-B datasets. Cluster densities are shown in light blue. The apo HKU1-A $S1^A$ structure (PDB 8OHN[3]) is shown as a black backbone trace in every panel as a common conformational reference. Percentages indicate the fraction of particles assigned to each cluster and sum to 100% for each dataset. The percentage shown in red marks the sole cluster whose density closely matches the apo HKU1-A $S1^A$ conformation.

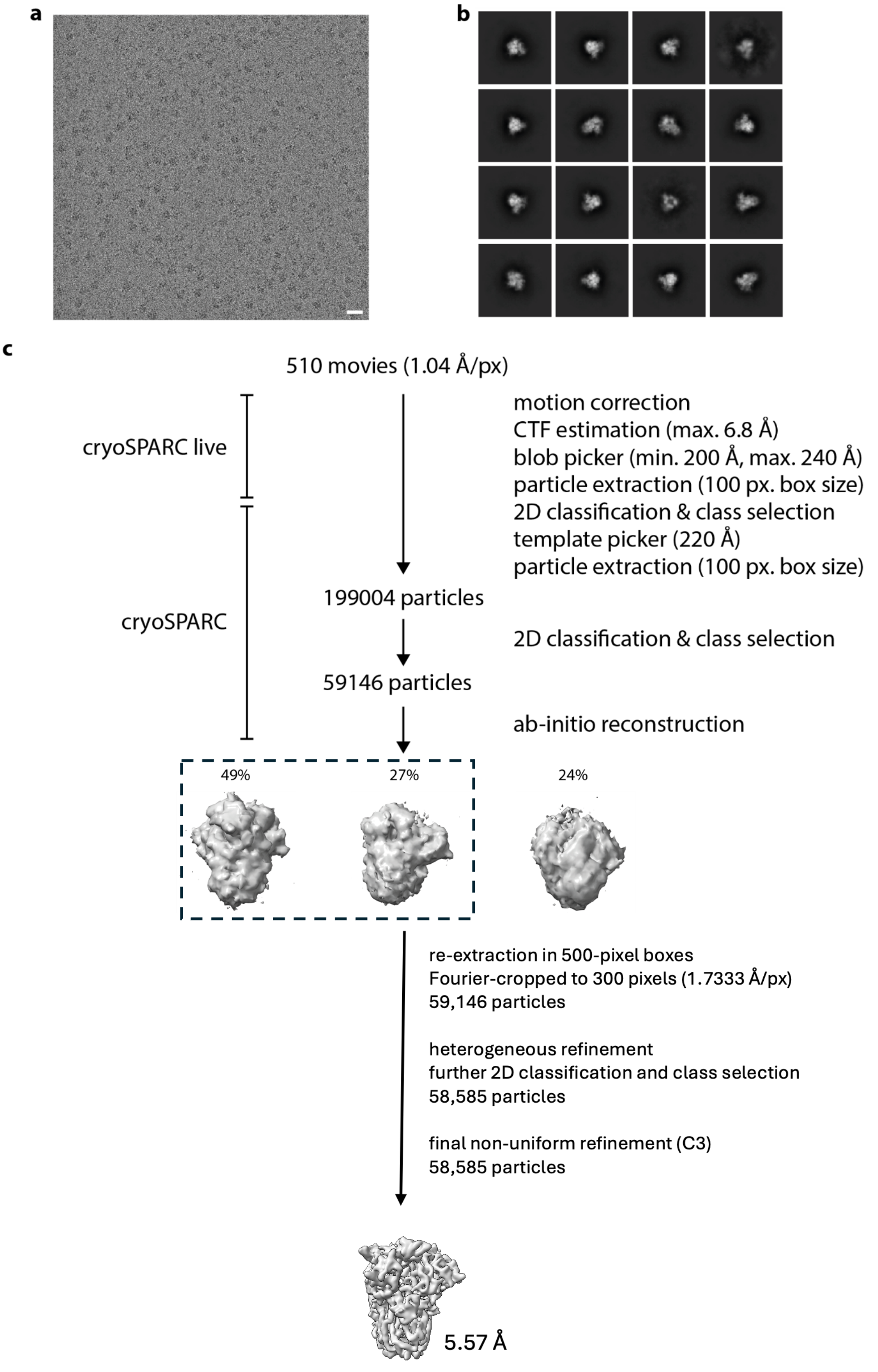


**Fig. S9: HKU1-B S proteins are closed after incubation with a disialoside ligand for 1 h at 37 °C.** **a.** Representative motion-corrected micrograph of S proteins collected on a 200-kV Talos Arctica microscope. The scale bar denotes 200 Å. **b.** Representative 2D class averages showing exclusively closed spike particles. **c.** Cryo-EM processing scheme through final C3 non-uniform refinement[1] of 58,585 particles, yielding a reconstruction at 5.57 Å resolution. No open spike class was detected at any stage of processing.

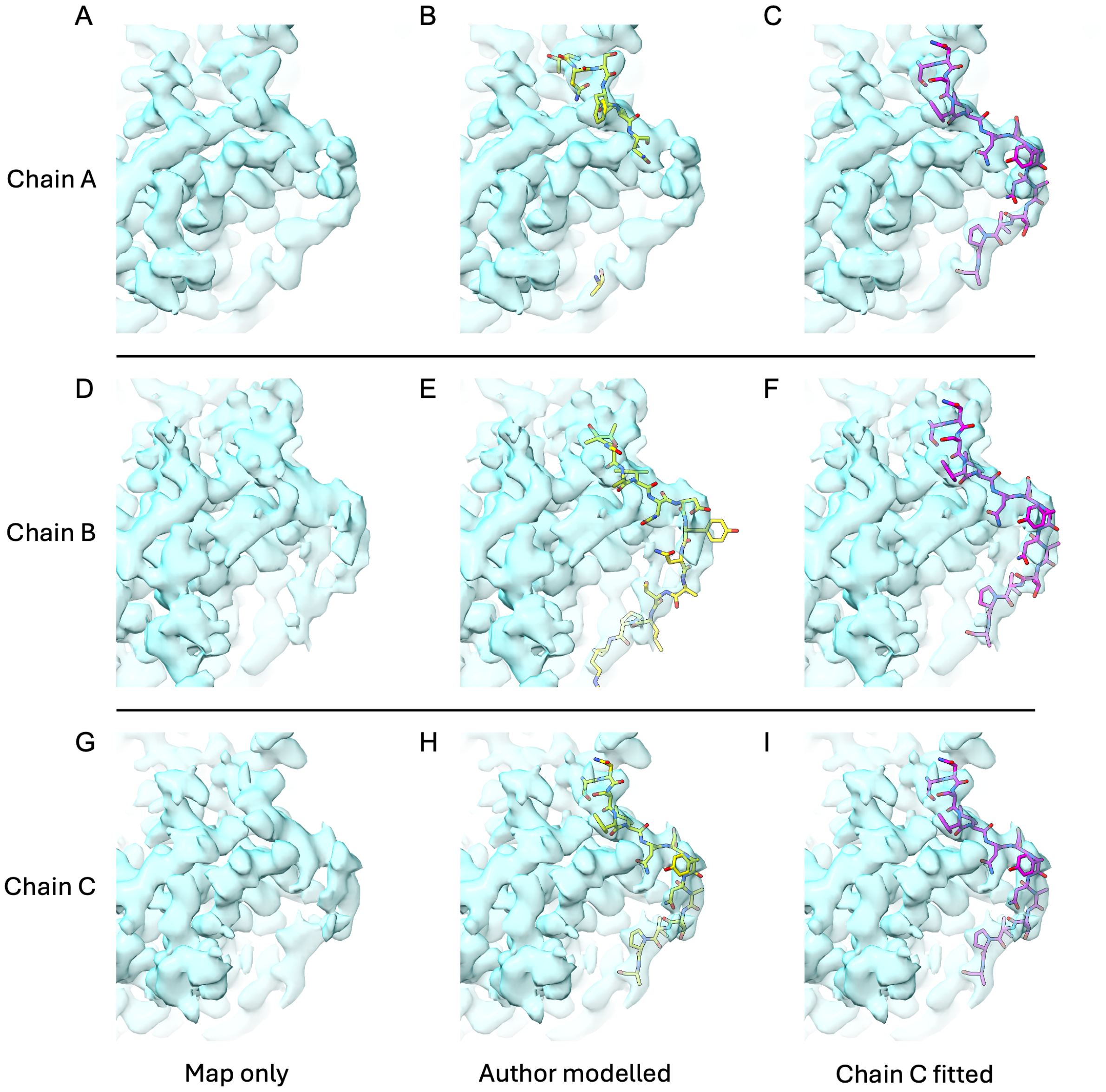


**Fig. S10: Reanalysis of a previously reported apo HKU1-B spike structure shows that all three e1-loop densities accommodate a common apo conformation.** Cryo-EM density corresponding to the e1 loop is shown for chains A (A–C), B (D–F) and C (G–I) of the apo HKU1-B spike reconstruction reported by Wang et al.[6] (EMDB-39041; PDB 8Y8C). Left panels show the experimental density alone. Middle panels show the deposited, author-modelled e1 conformations in yellow, which differ among the three chains. Right panels show the chain C e1-loop conformation, coloured magenta, fitted into the density of each chain. Despite the different conformations assigned in the deposited model, the density for all three protomers readily accommodates the same apo-like e1-loop conformation.

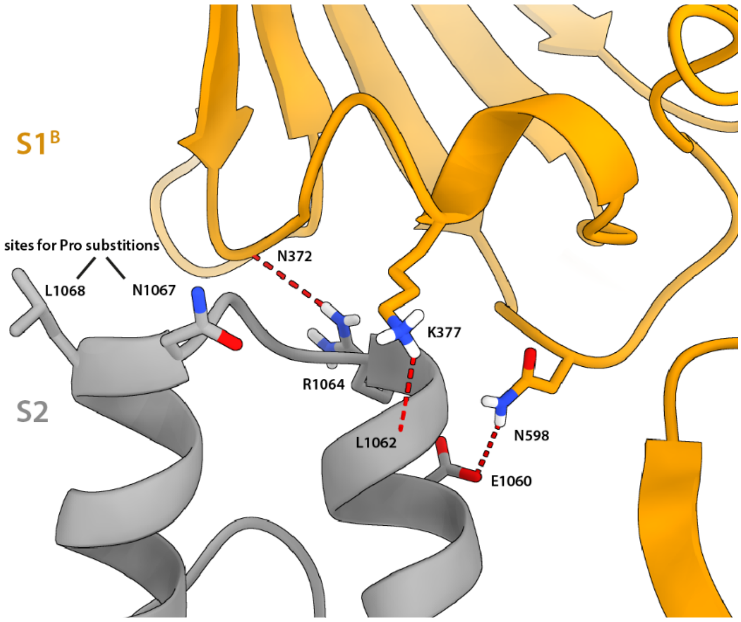


**Fig. S11: Interface between S2 and S1$^{B}$.** L1068 and N1067 in the helix-turn-helix element (gray) are commonly substituted by Pro residues to prevent S transition into the post-fusion state. However, these residues are part of an inter-domain interface with S1$^{B}$ (orange). H-bonds (red) that need to be broken for the transition into the S1$^{B}$-up state are indicated by dashed lines.

**a**

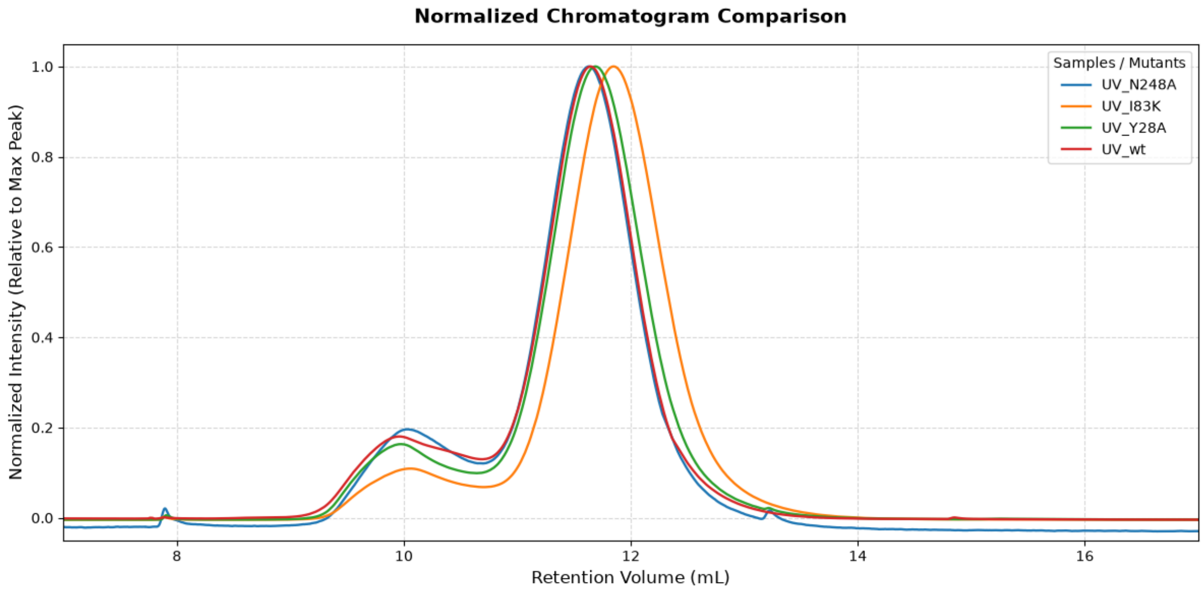


**b**

```
HKU1-B_N5        MFLIIFILPTTLAVIGDFNCTNSFINDYNKTIPRISEDVVDVSLGLGTYYVLNRVYLNTT 60
HKU1-B_N2        MFLIIFILPTTLAVIGDFNCTNSFINDYNKTIPRISEDVVDVSLGLGTYYVLNRVYLNTT 60
HKU1-A_Caen1     MLLIIFILPTTLAVIGDFNCTNFAINDLNTTIPRISEYVVDVSYGLGTYYILDRVYLNTT 60
HKU1-A_N1        MLLIIFILPTTLAVIGDFNCTNFAINDLNTTVPRISEYVVDVSYGLGTYYILDRVYLNTT 60
                 *:********************  *** *.*:***** ***** ******:*:*******

HKU1-B_N5        LLFTGYFPKSGANFRDLALKGSIYLSTLWYKPPFLSDFNNGIFSKVKNTKLYVNNTLYSE 120
HKU1-B_N2        LLFTGYFPKSGANFRDLALKGSKYLSTLWYKPPFLSDFNNGIFSKVKNTKLYVNNTLYSE 120
HKU1-A_Caen1     ILFTGYFPKSGANFRDLSLKGTTKLSTLWYQKPFLSDFNNGIFSRVKNTKLYVNKTLYSE 120
HKU1-A_N1        ILFTGYFPKSGANFRDLSLKGTTYLSTLWYQKPFLSDFNNGIFSRVKNTKLYVNKTLYSE 120
                 :****************:***:  ******: ************:*********:*****

HKU1-B_N5        FSTIVIGSVFVNTSYTIVVQPHNGILEITACQYTMCEYPHTVCKSKGSIRNESWHIDSSE 180
HKU1-B_N2        FSTIVIGSVFVNTSYTIVVQPHNGILEITACQYTMCEYPHTVCKSKGSIRNESWHIDSSE 180
HKU1-A_Caen1     FSTIVIGSVFINNSYTIVVQPHNGVLEITACQYTMCEYPHTICKSIGSSRNESWHFDKSE 180
HKU1-A_N1        FSTIVIGSVFINNSYTIVVQPHNGVLEITACQYTMCEYPHTICKSKGSSRNESWHFDKSE 180
                 **********:*.***********:****************:*** ** ******:*.**

HKU1-B_N5        PLCLFKKNFTYNVSADWLYFHFYQERGVFYAYYADVGMPTTFLFSLYLGTILSHYYVMPL 240
HKU1-B_N2        PLCLFKKNFTYNVSADWLYFHFYQERGVFYAYYADVGMPTTFLFSLYLGTILSHYYVMPL 240
HKU1-A_Caen1     PLCLFKKNFTYNVSTDWLYFHFYQERGTFYAYYADSGMPTTFLFSLYLGTLLSHYYVLPL 240
HKU1-A_N1        PLCLFKKNFTYNVSTDWLYFHFYQERGTFYAYYADSGMPTTFLFSLYLGTLLSHYYVLPL 240
                 **************:************.******* **************:******:**

HKU1-B_N5        TCNAISSNTDNETLEYWVTPLSRRQYLLNFDEHGVITNAVDCSSSFLSEIQCKTQSFAPN 300
HKU1-B_N2        TCKAISSNTDNETLEYWVTPLSRRQYLLNFDEHGVITNAVDCSSSFLSEIQCKTQSFAPN 300
HKU1-A_Caen1     TCNAISSNTDNETLQYWVTPLSKRQYLLKFDDRGVITNAVDCSSSFFSEIQCKTKSLLPN 300
HKU1-A_N1        TCNAISSNTDNETLQYWVTPLSKRQYLLKFDNRGVITNAVDCSSSFFSEIQCKTKSLLPN 300
                 **:***********:*******:*****:**::*************:*******:*:*: **
```

**Fig. S12: Normalized SEC data from HKU1-B S1$^{A}$ hFc mutant proteins used for BLI studies (a) and multiple sequence alignment of S1$^{A}$ domains from selected isolates of HKU1-A and -B.** Protein identifiers are as follows: ADN03339.1 (Caen1, GenBank), AAT98580.1 (N1, GenBank), Q0ZME7 (N5, UniProt), Q14EB0 (N2, UniProt). Alignments were done using the clustalo program v1.2.4[7].

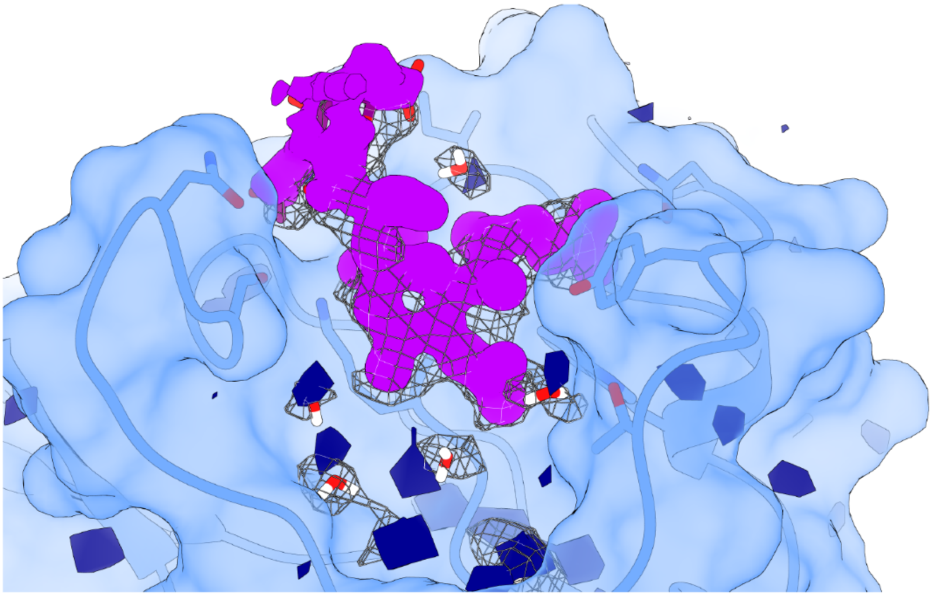

**Fig. S13: Comparison of experimental cryo-EM density (mesh) and MD-derived atomic density of water molecules (dark blue) and the disialoside ligand (pink).** MD simulation (100 ns, backbone atoms restrained) was performed using the experimental structure of $S1^{A}$ (residues 14–298) with $Man_5$ N-glycans attached.

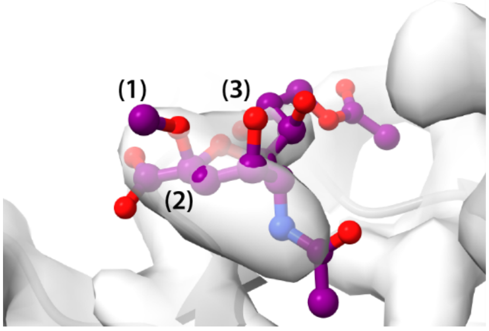


**Fig. S14: Sialic acid model in PDB 8Y8J**. The structure modeled by Wang et al.[6] depicts an α-methylglycoside (**1**) despite the fact that unmodified sialic acid was reportedly added to the sample which would predominantly adopt a β-anomeric configuration in solution[8]. Furthermore, the modelled bond angle at C2 is 90° (2), substantially deviating from the natural geometry of $sp^3$ hybridized carbon atoms. Lastly, sialic acid is shown in an unusual, high-energy boat conformation[8] (**3**), with O4 clearly outside of the experimental density (gray surface, EMD-39048).

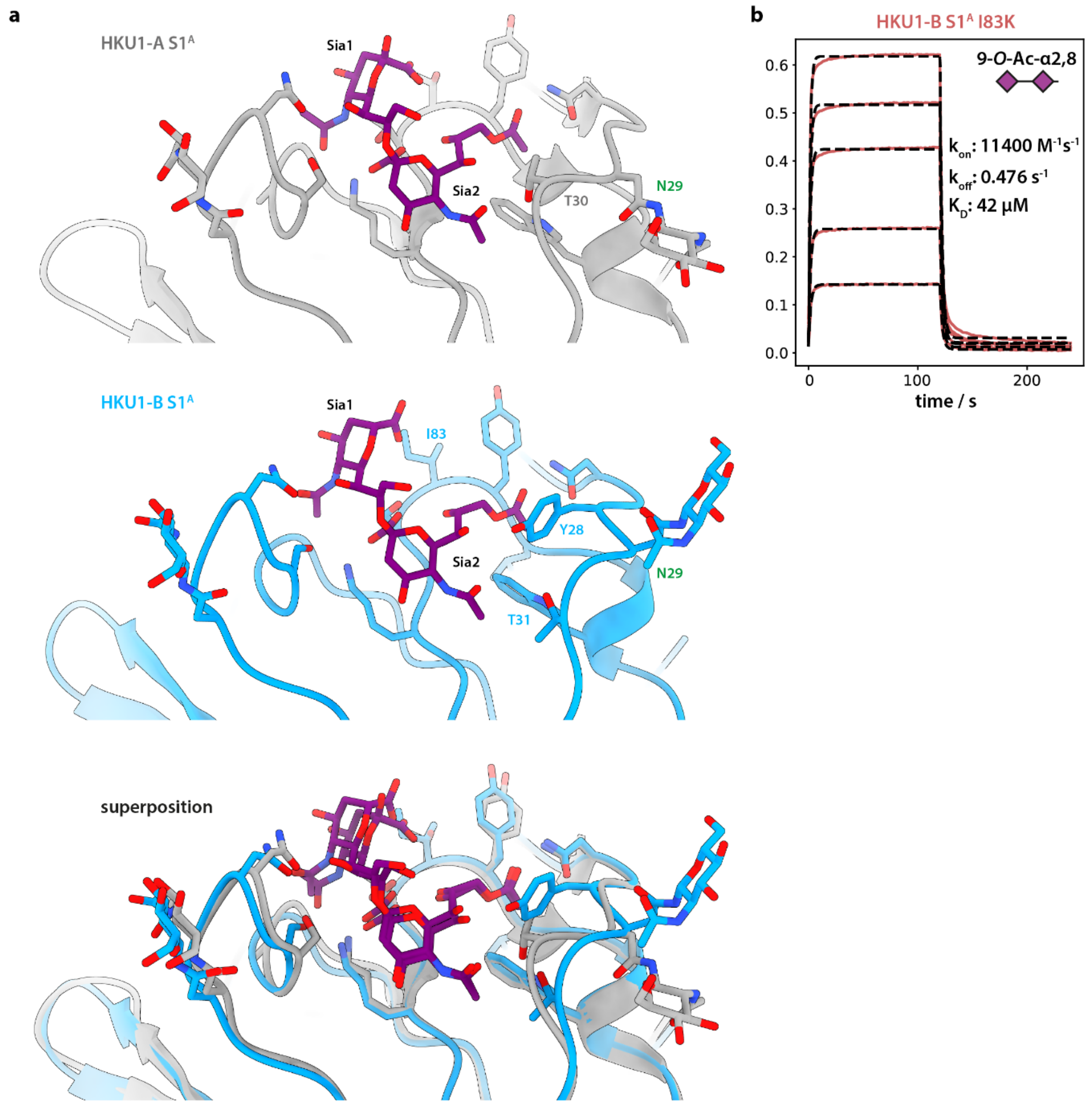


**Fig. S15**: **The extended Sia binding site in HKU1-A and -B S1$^A$. a.** Binding site topology in HKU1-A and -B S1$^A$ domains as found in PDB 9BTB[9] (grey) and the structure determined in this study (blue). The disialoside binding pocket is generally conserved among both HKU1 serotypes, showing similar binding poses of the terminal Sia2 and the penultimate Sia1. The interaction with Tyr28, unique for B-type S, seems to cause local conformational changes in e1, including an altered conformation of the N29 glycan and different hydrogen bond patterns around Thr30/31, respectively. **b.** Natural HKU1-B variants with K at position 83 show enhanced binding as demonstrated by analysis of kinetic BLI data of I83K S1$^A$ monomers, presumably by electrostatic interactions with the carboxylate group of Sia1.

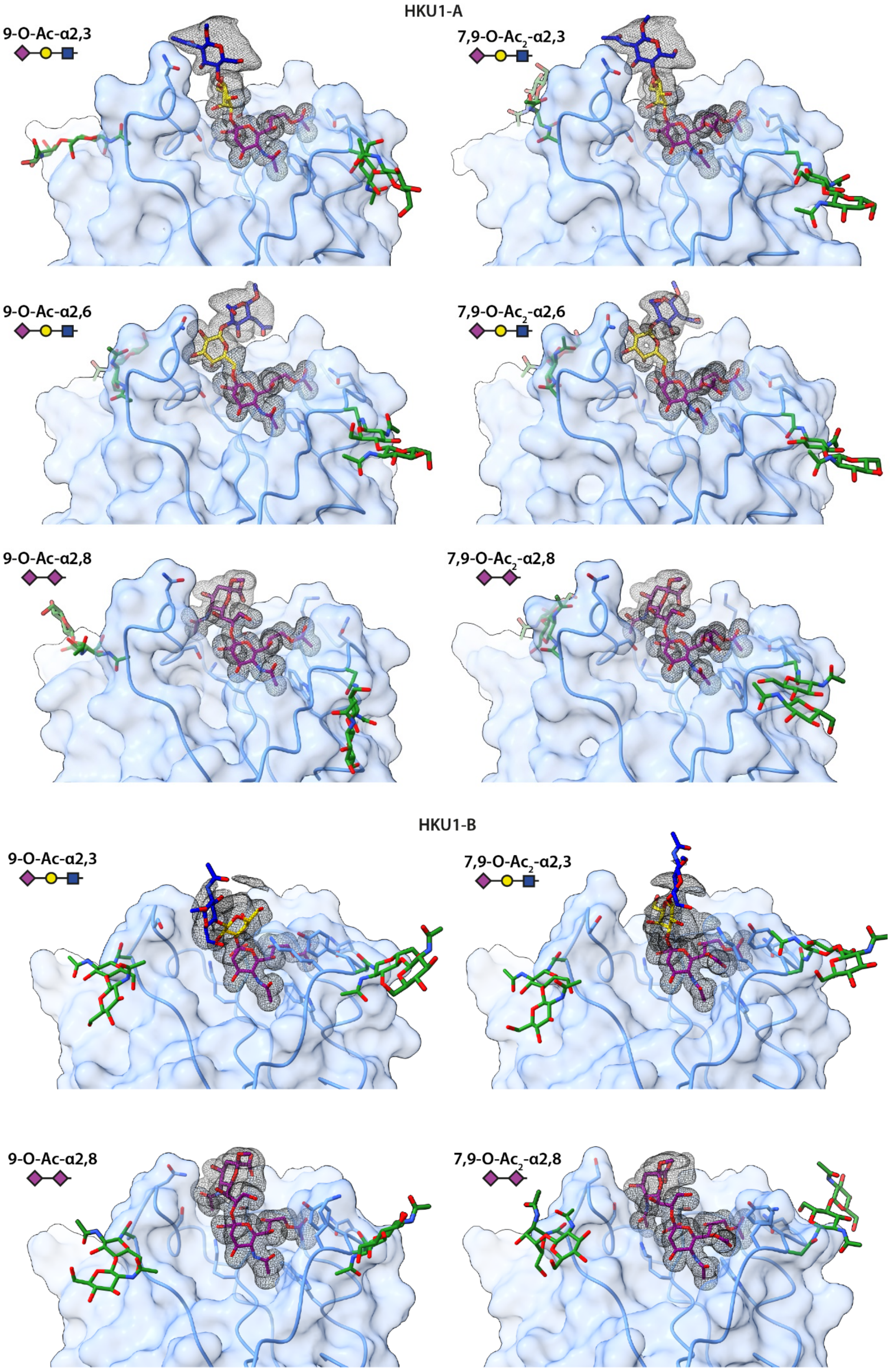
HKU1-A
9-O-Ac-α2,3
7,9-O-Ac2-α2,3
9-O-Ac-α2,6
7,9-O-Ac2-α2,6
9-O-Ac-α2,8
7,9-O-Ac2-α2,8
HKU1-B
9-O-Ac-α2,3
7,9-O-Ac2-α2,3
9-O-Ac-α2,8
7,9-O-Ac2-α2,8

**Fig. S16: Molecular dynamics-derived binding poses of different 9-O-Ac sialosides.** Average atomic densities over the course of the respective 300 ns simulations are shown as meshes and the best fitting single glycan structure was selected for display. Blurriness is indicative of ligand flexibility. For 2,3- and 2,6-linked glycans, relevant contacts are only made via the terminal Sia2. Only the 2,8-linked disialoside shows additional interactions with the extended binding site in e2. PDB 8OPM[3] was the basis for the depicted HKU1-A binding site. Simulations were performed using $S1^{A}$ (residues 14–298) with $Man_5$ N-glycans attached.

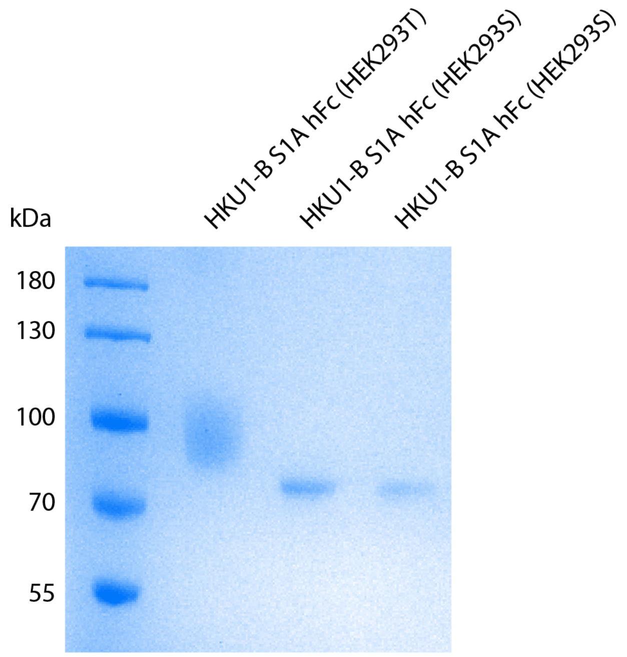


**Fig. S17: HKU1-A and -B S1$^{A}$ hFc proteins show reduced, more uniform glycosylation when expressed in HEK293S cells.** A 10% SDS-PAGE gel of samples used for BLI experiments shows the mobility shift and band narrowing expected from uniform $Man_5$-glycosylation in GnTI-deficient HEK293S cells.

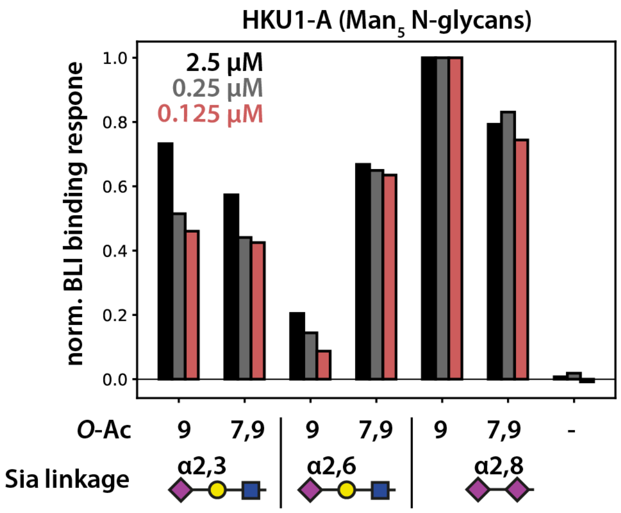


**Fig. S18: BLI binding levels at different probe concentrations of $Man_5$-glycosylated HKU1-A $S1^A$ Fc proteins, expressed in HEK293S cells.** Data were normalized to the respective binding response to the 9-*O*-Ac disialoside.

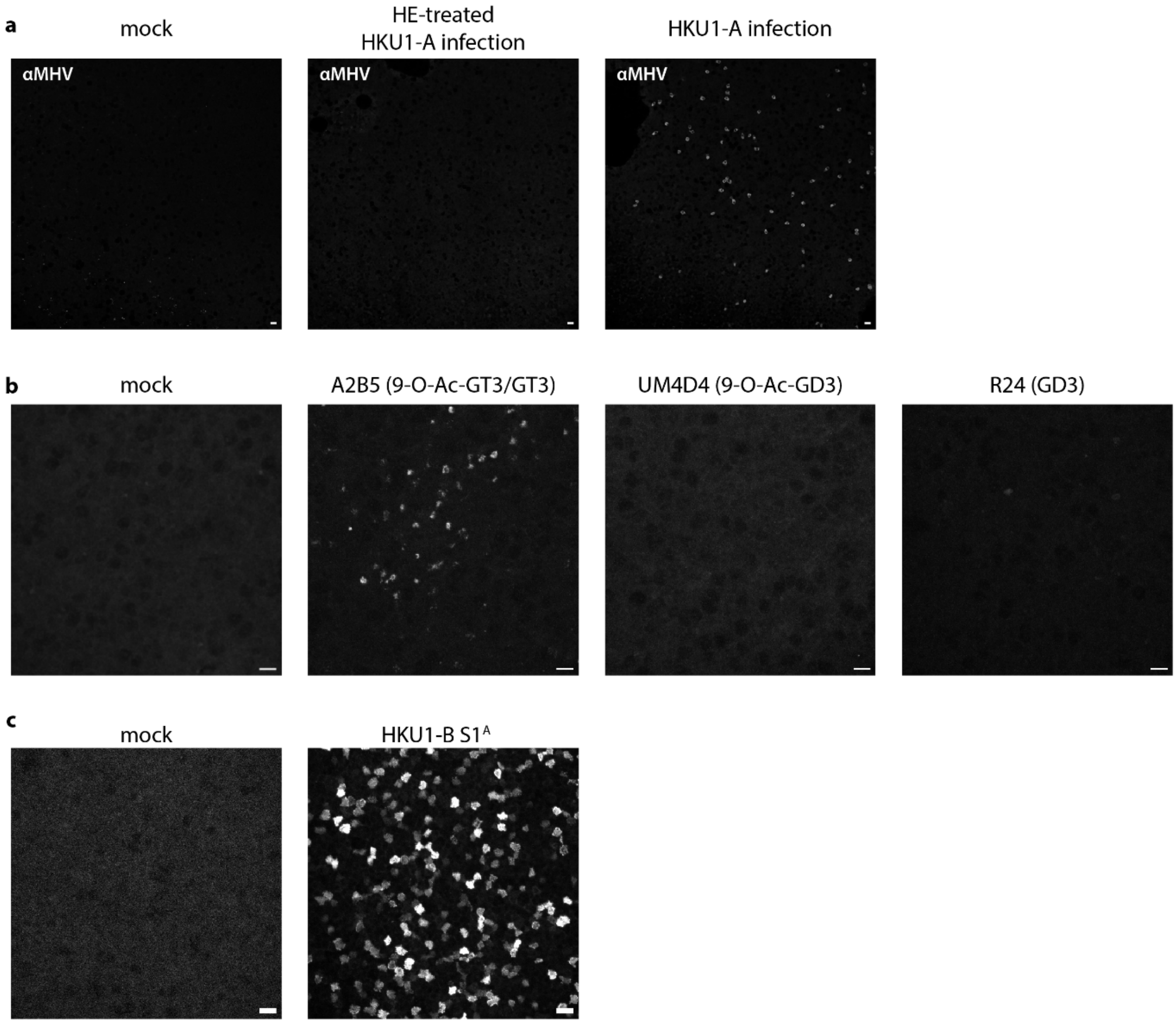


**Fig. S19: Anti-ganglioside staining experiments and control experiments under mock conditions.** **a.** αMHV-staining experiments 24 h after infection with HKU1-A Caen1. Images are stitched overview images and were used for quantification of infection levels as shown in Fig. 4a, b. **b.** Anti-ganglioside antibodies reveal the presence of GT3-like gangliosides but not GD3 in human nasal epithelial cells. Cell surfaces of non-permeabilized cells present A2B5 but not R24 or UM4D4 glycotopes. **c.** HKU1-B $S1^A$ staining experiment as compared to the negative control. All scale bars denote 20 µm. Images in b and c are maximum intensity projections of confocal stacks.

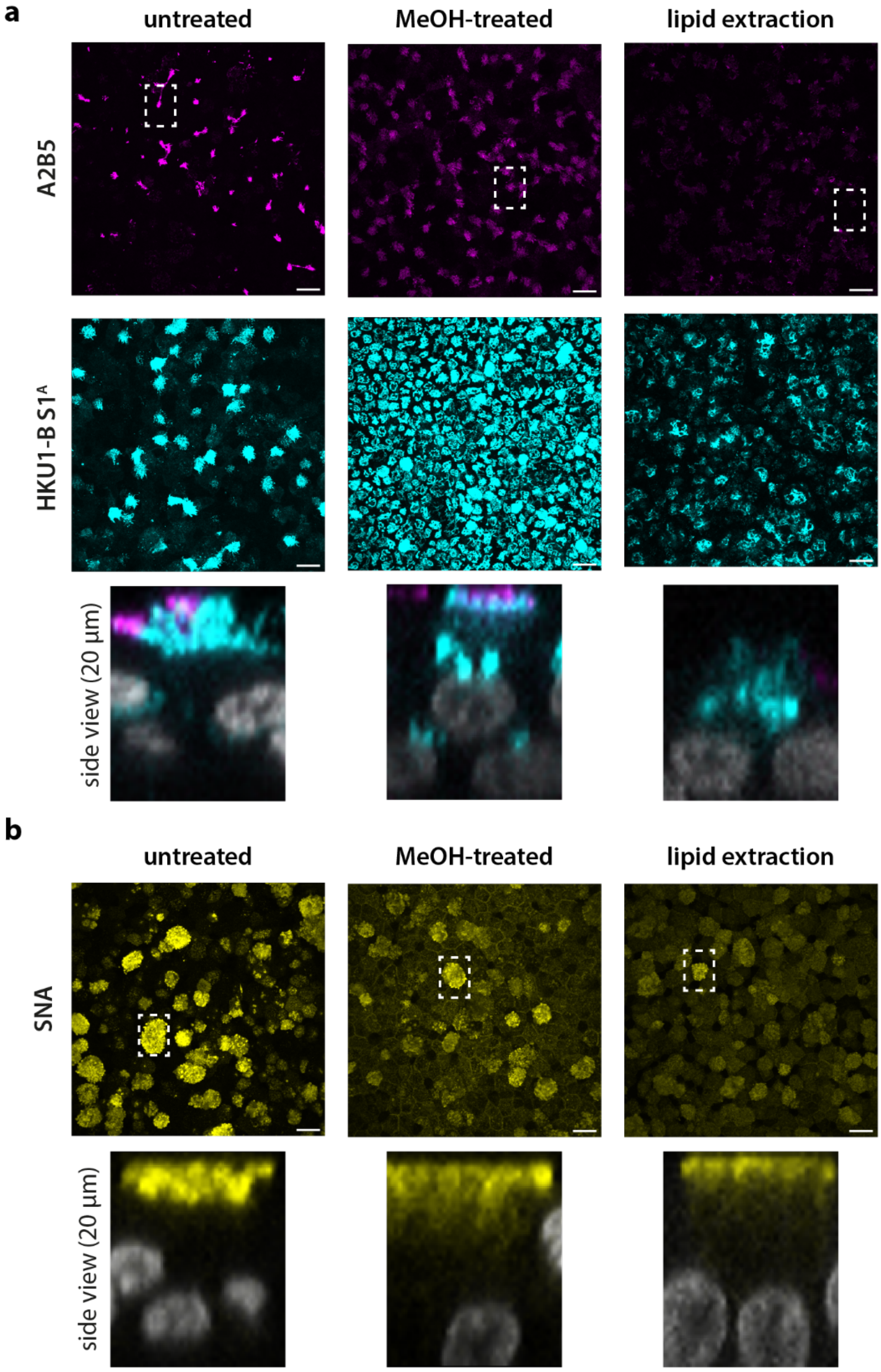


**Fig. S20: A2B5 staining is sensitive to methanol treatment and lipid extraction. a.** Methanol (MeOH) treatment of HNECs decreases fluorescence intensity of antibody A2B5. Subjecting cells to a series of ethanol and xylene solutions almost completely abolishes A2B5 staining, indicating that extractable glycolipids are the main source of A2B5 staining. HKU1-B S1$^{A}$ surface staining is restricted to a subpopulation of cells but permeabilization with MeOH yields indiscriminate, perinuclear staining. Surface staining of HKU1-B S1$^{A}$ is mostly lost after lipid extraction, but perinuclear staining is more resistant. **b.** SNA lectin was used as a control, primarily recognizing surface-bound 2,6-linked sialoglycoproteins. Scale bars denote 20 µm. Top view images are maximum intensity projections of confocal stacks. Side views are slices through the confocal stacks in the indicated regions of the overview images.

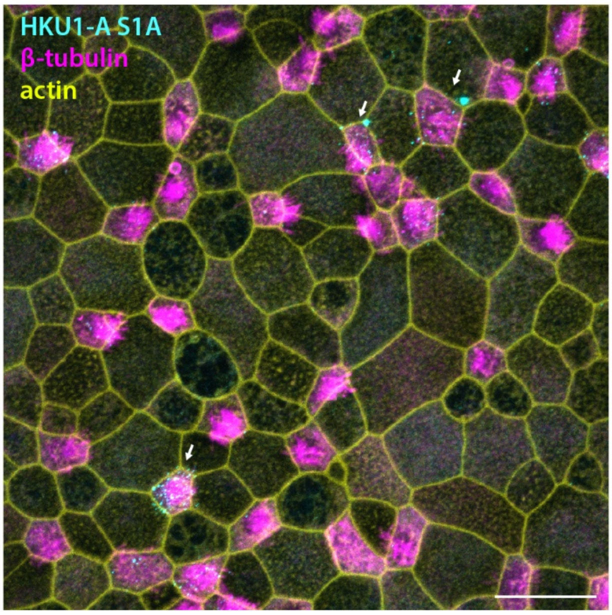


**Fig. S21: HKU1-A S1$^{A}$ domains recognize ciliated cells in the nasal epithelium.** Surface immunofluorescence staining with HKU1-A S1$^{A}$ domains results in weaker fluorescence than comparable experiments with HKU1-B (Fig. 4). β-tubulin staining (magenta) was used as a marker for ciliated cells.

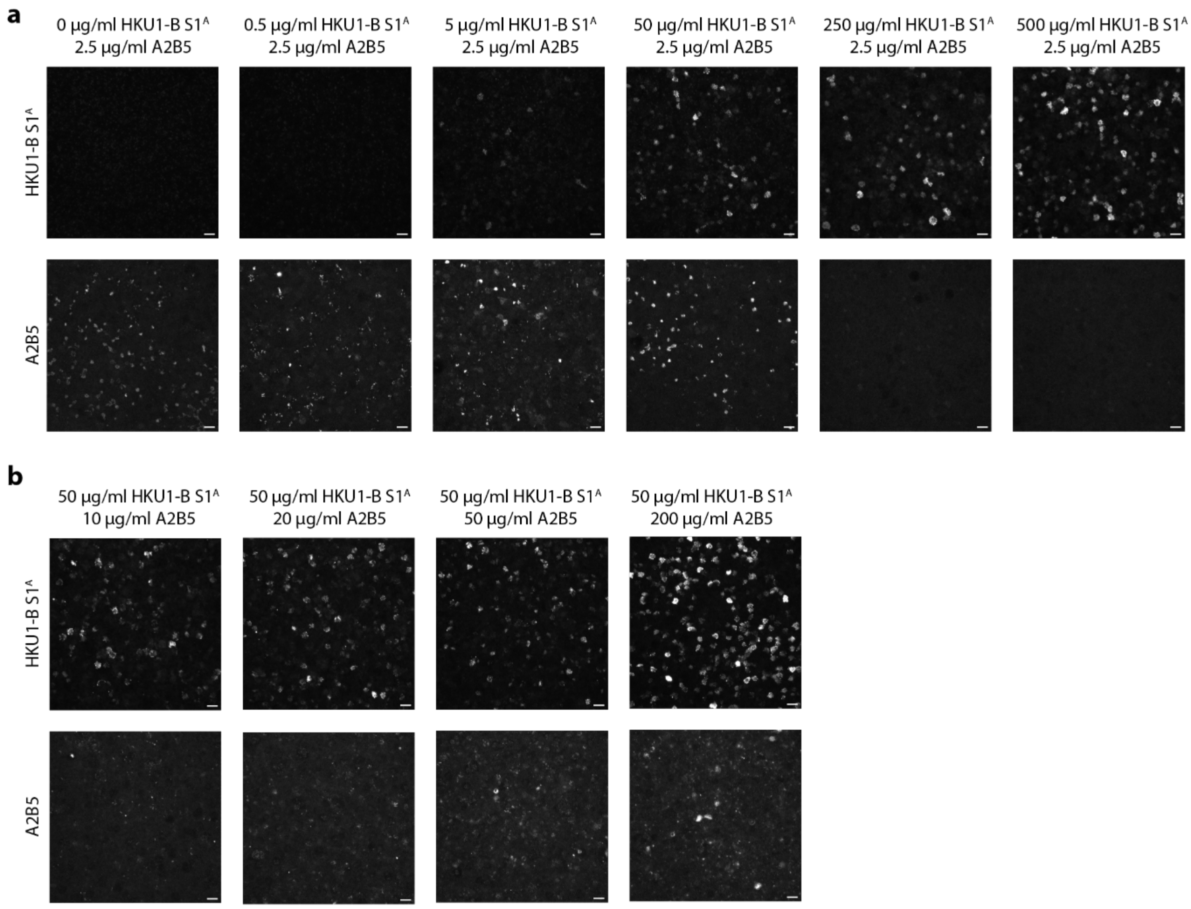


**Fig. S22: HKU1 S1$^{A}$ domains compete with A2B5 antibodies for binding of cell surface sialoglycans. a.** Increasing concentrations of HKU1-B S1$^{A}$ proteins prohibit binding of antibody A2B5. **b.** Conversely, increasing A2B5 protein concentration does not influence staining intensity of HKU1 S1$^{A}$. Scale bars denote 20 µm. Images are maximum intensity projections of confocal stacks.

**a**

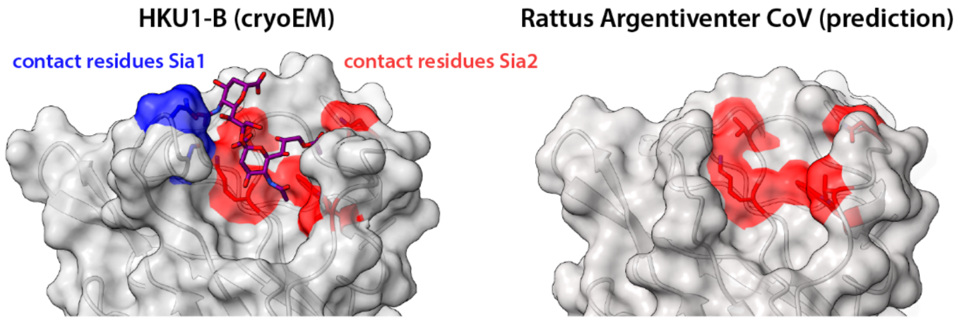


**b**

```
N5     14     VIGDFNCTNSFINDYNKTIPRISEDVVDVSLGLGTYYVLNRVYLNTTLLFTGYFPKSGAN  73
RCoV   14     VIGDFKCTTLSINDVNTGVPSISSEVVDVSNGLGTYYVLDRVYLNTTLLLNGYYPISGAS  73

N5     74     FRDLALKGSIYLSTLWYKPPFLSDFNNGIFSKVKNTKLYVNNTLYSEFSTIVIGSVFVNT  133
RCoV   74     FRNLALKGTLLLSTLWYKPPFLSEFNNGIFAKVRNTKLITNGVTYSEFNTIVIGSVFVNT  133

N5     134    SYTIVVQPHNGILEITACQYTMCEYPHTVCKSK-GSIRNESWHID-SSEPLCLFKKNFTY  191
RCoV   134    SYTIVVQPHNGILEITACQYTMCEYPNTNCHSALGNQRQELWHYEGNNQPLCLFKKNFTY  193

N5     192    NVSADWLYFHFYQERGVFYAYYADVGMPTTFLFSLYLGTILSHYYVMPLTCNAISSNTDN  251
RCoV   194    DVNVDWLYFHFYQEGGTFYAYYADHSLPTTFLFSLYIGSVLSHYYVVPLTCNAQAPN---  250

N5     252    ETLEYWVTPLSRRQYLLNFDEHGVITNAVDCSSSFLSEIQCKTQSFAPNTGVYDLSGFTV  311
RCoV   251    --MQYWVTPLAKRQYLLKFDERGVITNAVDCVSNYLSEIQCKTQSLTPNTGVYDLSGYTV  308
```

**Fig. S23: Predicted structure of the S1$^{A}$ domain of *Rattus argentiventer* CoV (RCoV), the closest known murine relative of HKU1. a.** *Rattus argentiventer* CoV (GenBank: AYR18616.1) was identified as HKU1's closest relative in a BLAST search[10] using HKU1-B S as query and filtering for non-human hosts. Comparing the experimental structure of HKU1-B S with the AlphaFold 3 model[11] (as accessed through the AlphaFold webserver in May 2025) of RCoV reveals the structural conservation of key contact residues of Sia2, suggesting an ancestral binding site for 9-*O*-Ac sialosides (red residues). Residues in contact with Sia1 in HKU1 S (blue residues), however, are absent in RCoV due to the missing e2 loop. **b**. Sequence alignment of RCoV S1$^{A}$ with the HKU1-B N5 reference sequence (Q0ZME7 UniProt).

**Table S1. Cryo-EM data collection, refinement and validation statistics for global and local refinements.**

| | HKU1-B apo Global refinement (EMD-XXXX; PDB XXXX) | HKU1-B holo Global refinement (EMD-XXXX; PDB XXXX) | HKU1-B holo S1A local refinement (EMD-XXXX) | HKU1-B holo, 37 °C Global refinement (EMD-XXXX) |
|---|---|---|---|---|
| **Data collection and processing** | | | | |
| Microscope | Titan Krios G4 | Titan Krios G4 | Titan Krios G4 | Talos Arctica |
| Detector | Falcon 4i | Falcon 4i | Falcon 4i | K2 Summit |
| Energy filter (slit width) | Selectris X (10 eV) | Selectris X (10 eV) | Selectris X (10 eV) | GIF Quantum (20 eV) |
| Acceleration voltage (kV) | 300 | 300 | 300 | 200 |
| Recording mode | EER | EER | EER | Counting |
| Nominal magnification | 165,000× | 165,000× | 165,000× | 130,000× |
| Calibrated pixel size (Å) | 0.73 | 0.73 | 0.73 | 1.04 |
| Stage tilt (°) | 32 | 32 | 32 | 32 |
| Exposure time (s) | 3.46 | 3.46 | 3.46 | 10.0 |
| Electron exposure ($e^- Å^{-2}$) | 50 | 50 | 50 | 50 |
| Defocus range (µm) | −0.5 to −2.0 | −0.5 to −2.0 | −0.5 to −2.0 | −0.5 to −2.0 |
| Movies collected (no.) | 2,445 | 5,402 | 5,402 | 510 |
| Symmetry imposed | C3 | C3 | C1 | C3 |
| Particle images (no.) | 61,233 | 196,925 | 590,775 | 58,585 |
| Reconstruction box / deposited voxel size | 500 px / 0.73 Å | 500 px / 0.73 Å | 500 px / 0.73 Å | 300 px / 1.7333 Å |
| Map resolution (Å), FSC 0.143 | 2.57 | 2.40 | 2.74 | 5.57 |
| Local resolution range (Å), FSC 0.5 (minimum–maximum) | 2.25–42.77 | 2.01–37.83 | 2.30–9.68 | 5.02–12.93 |
| Map sharpening B factor (Å²) | −63.06 | −71.45 | −101.33 | −398.3 |
| **Refinement** | | | | |
| Initial model used | PDB 5I08 | PDB 5I08 | — | — |
| Model resolution (Å), FSC 0.5 | 2.70 | 2.46 | — | — |
| Model-to-map CC (mask / volume) | 0.8321 / 0.7998 | 0.8035 / 0.7698 | — | — |
| **Model composition** | | | | |
| Non-hydrogen atoms | 28,779 | 29,058 | — | — |
| Protein residues | 3,525 | 3,534 | — | — |
| Ligands | 90 | 99 | — | — |
| Solvent molecules | 0 | 21 | — | — |
| **Mean B factors (Å²)** | | | | |
| Protein | 44.69 | 33.74 | — | — |
| Ligands | 97.90 | 76.27 | — | — |
| Solvent | — | 57.48 | — | — |
| **R.m.s. deviations** | | | | |
| Bond lengths (Å) | 0.003 | 0.008 | — | — |
| Bond angles (°) | 0.620 | 1.014 | — | — |
| **Validation** | | | | |
| MolProbity score | *0.90* | *0.88* | — | — |
| Clashscore | 1.35 | 0.99 | — | — |
| Poor rotamers (%) | 0.00 | 0.00 | — | — |
| **Ramachandran plot** | | | | |
| Favoured (%) | 97.85 | 97.60 | — | — |
| Allowed (%) | 2.15 | 2.40 | — | — |
| Outliers (%) | 0.00 | 0.00 | — | — |

**Table S2. Amino acid sequences of synthesized proteins.** Thrombin cleavage sites are indicated in red for relevant constructs.

| | |
|---|---|
| HKU1-A S1[A] hFc | VIGDFNCTNFAINDLNTTIPRISEYVVDVSYGLGTYYILDRVYLNTTILFTGYFPKSGANFRDLSLKGTTKLSTLWYQ<br>KPFLSDFNNGIFSRVKNTKLYVNKTLYSEFSTIVIGSVFINNSYTIVVQPHNGVLEITACQYTMCEYPHTICKSIGSS<br>RNESWHFDKSEPLCLFKKNFTYNVSTDWLYFHFYQERGTFYAYYADSGMPTTFLFSLYLGTLLSHYYVLPLTCNAISS<br>NTDNETLQYWVTPLSKRQYLLKFDDRGVITNAVDCSSSFFSEIQCKTKSLLDPLVPRGSGGGGDPEPKSCDKTHTCPP<br>CPAPELLGGPSVFLFPPKPKDTLMISRTPEVTCVVVDVSHEDPEVKFNWYVDGVEVHNAKTKPREEQYNSTYRVVSVL<br>TVLHQDWLNGKEYKCKVSNKALPAPIEKTISKAKGQPREPQVYTLPPSRDELTKNQVSLTCLVKGFYPSDIAVEWESN<br>GQPENNYKTTPPVLDSDGSFFLYSKLTVDKSRWQQGNVFSCSVMHEALHNHYTQKSLSLSPGK |
| HKU1-B S1[A] hFc | VIGDFNCTNSFINDYNKTIPRISEDVVDVSLGLGTYYVLNRVYLNTTLLFTGYFPKSGANFRDLALKGSIYLSTLWYK<br>PPFLSDFNNGIFSKVKNTKLYVNNTLYSEFSTIVIGSVFVNTSYTIVVQPHNGILEITACQYTMCEYPHTVCKSKGSI<br>RNESWHIDSSEPLCLFKKNFTYNVSADWLYFHFYQERGVFYAYYADVGMPTTFLFSLYLGTILSHYYVMPLTCNAISS<br>NTDNETLEYWVTPLSRRQYLLNFDEHGVITNAVDCSSSFLSEIQCKTQSFADPLVPRGSGGGGDPEPKSCDKTHTCPP<br>CPAPELLGGPSVFLFPPKPKDTLMISRTPEVTCVVVDVSHEDPEVKFNWYVDGVEVHNAKTKPREEQYNSTYRVVSVL<br>TVLHQDWLNGKEYKCKVSNKALPAPIEKTISKAKGQPREPQVYTLPPSRDELTKNQVSLTCLVKGFYPSDIAVEWESN<br>GQPENNYKTTPPVLDSDGSFFLYSKLTVDKSRWQQGNVFSCSVMHEALHNHYTQKSLSLSPGK |
| HKU1-A ecto | VIGDFNCTNFAINDLNTTIPRISEYVVDVSYGLGTYYILDRVYLNTTILFTGYFPKSGANFRDLSLKGTTKLSTLWYQ<br>KPFLSDFNNGIFSRVKNTKLYVNKTLYSEFSTIVIGSVFINNSYTIVVQPHNGVLEITACQYTMCEYPHTICKSIGSS<br>RNESWHFDKSEPLCLFKKNFTYNVSTDWLYFHFYQERGTFYAYYADSGMPTTFLFSLYLGTLLSHYYVLPLTCNAISS<br>NTDNETLQYWVTPLSKRQYLLKFDDRGVITNAVDCSSSFFSEIQCKTKSLLPNTGVYDLSGFTVKPVATVHRRIPDLP<br>DCDIDKWLNNFNVPSPLNWERKIFSNCNFNLSTLLRLVHTDSFSCNNFDESKIYGSCFKSIVLDKFAIPNSRRSDLQL<br>GSSGFLQSSNYKIDTTSSSCQLYYSLPAINVTINNYNPSSWNRRYGFNNFNLSSHSVVYSRYCFSVNNTFCPCAKPSF<br>ASSCKSHKPPSASCPIGTNYRSCESTTVLDHTDWCRCSCLPDPITAYDPRSCSQKKSLVGVGEHCAGFGVDEEKCGVL<br>DGSYNVSCLCSTDAFLGWSYDTCVSNNRCNIFSNFILNGINSGTTCSNDLLQPNTEVFTDVCVDYDLYGITGQGIFKE<br>VSAVYYNSWQNLLYDFNGNIIGFKDFVTNKTYNIFPCYAGRVSAAFHQNASSLALLYRNLKCSYVLNNISLATQPYFD<br>SYLGCVFNADNLTDYSVSSCALRMGSGFCVDYNSPSSSSSGGSGSSISASYRFVTFEPFNVSFVNDSIESVGGLYEIK<br>IPTNFTIVGQEEFIQTNSPKVTIDCSLFVCSNYAACHDLLSEYGTFCDNINSILDEVNGLLDTTQLHVADTLMQGVTL<br>SSNLNTNLHFDVDNINFKSLVGCLGPHCGSSSRSFFEDLLFDKVKLSDVGFVEAYNNCTGGSEIRDLLCVQSFNGIKV<br>LPPILSESQISGYTTAATVAAMFPPWSAAAGIPFSLNVQYRINGLGVTMDVLNKNQKLIATAFNNALLSIQNGFSATN<br>SALAKIQSVVNSNAQALNSLLQQLFNKFGAISSSLQEILSRLDALEAQVQIDRLINGRLTALNAYVSQQLSDISLVKL<br>GAALAMEKVNECVKSQSPRINFCGNGNHILSLVQNAPYGLLFMHFSYKPISFKTVLVSPGLCISGDVGIAPKQGYFIK<br>HNDHWMFTGSSYYYPEPISDKNVVFMNTCSVNFTKAPLVYLNHSVPKLSDFESELSHWFKNQTSIAPNLTLNLHTINA<br>TFLDLLIKRMKQIEDKIEEIESKQKKIENEIARIKKIKLVPRGSLEWSHPQFEK |
| BCoV HE hFc | FDNPPTNVVSHLNGDWFLFGDSRSDCNHVVNTNPRNYSYMDLNPALCDSGKISSKAGNSIFRSFHFTDFYNYTGEGQQ<br>IIFYEGVNFTPYHAFKCTTSGSNDIWMQNKGLFYTQVYKNMAVYRSLTFVNVPYVYNGSAQSTALCKSGSLVLNNPAY<br>IAREANFGDYYYKVEADFYLSGCDEYIVPLCIFNGKFLSNTKYYDDSQYYFNKDTGVIYGLNSTETITTGFDFNCHYL<br>VLPSGNYLAISNELLLTVPTKAICLNKRKDFTPVQVVDSRWNNARQSDNMTAVACQPPYCYFRNSTTNYVGVYDINHG<br>DAGFTSILSGLLYDSPCFSQQGVFRYDNVSSVWPLYSYGRCPTAADINTPDVPICVYDSDPLVPRGSDPEPKSCDKTH<br>TCPPCPAPELLGGPSVFLFPPKPKDTLMISRTPEVTCVVVDVSHEDPEVKFNWYVDGVEVHNAKTKPREEQYNSTYRV<br>VSVLTVLHQDWLNGKEYKCKVSNKALPAPIEKTISKAKGQPREPQVYTLPPSRDELTKNQVSLTCLVKGFYPSDIAVE<br>WESNGQPENNYKTTPPVLDSDGSFFLYSKLTVDKSRWQQGNVFSCSVMHEALHNHYTQKSLSLSPGK |
| HKU1-B ecto | MPMGSLQPLATLYLLGMLVASVLAVIGDFNCTNSFINDYNKTIPRISEDVVDVSLGLGTYYVLNRVYLNTTLLFTGYF<br>PKSGANFRDLALKGSIYLSTLWYKPPFLSDFNNGIFSKVKNTKLYVNNTLYSEFSTIVIGSVFVNTSYTIVVQPHNGI<br>LEITACQYTMCEYPHTVCKSKGSIRNESWHIDSSEPLCLFKKNFTYNVSADWLYFHFYQERGVFYAYYADVGMPTTFL<br>FSLYLGTILSHYYVMPLTCNAISSNTDNETLEYWVTPLSRRQYLLNFDEHGVITNAVDCSSSFLSEIQCKTQSFAPNT<br>GVYDLSGFTVKPVATVYRRIPNLPDCDIDNWLNNVSVPSPLNWERRIFSNCNFNLSTLLRLVHVDSFSCNNLDKSKIF<br>GSCFNSITVDKFAIPNRRRDDLQLGSSGFLQSSNYKIDISSSSCQLYYSLPLVNVTINNFNPSSWNRRYGFGSFNLSS<br>YDVVYSDHCFSVNSDFCPCADPSVVNSCAKSKPPSAICPAGTKYRHCDLDTTLYVKNWCRCSCLPDPISTYSPNTCPQ<br>KKVVVGIGEHCPGLGINEEKCGTQLNHSSCFCSPDAFLGWSFDSCISNNRCNIFSNFIFNGINSGTTCSNDLLYSNTE<br>ISTGVCVNYDLYGITGQGIFKEVSAAYYNNWQNLLYDSNGNIIGFKDFLTNKTYTILPCYSGRVSAAFYQNSSSPALL<br>YRNLKCSYVLNNISFISQPFYFDSYLGCVLNAVNLTSYSVSSCDLRMGSGFCIDYALPSSGGSGSGISSPYRFVTFEP<br>FNVSFVNDSVETVGGLFEIQIPTNFTIAGHEEFIQTSSPKVTIDCSAFVCSNYAACHDLLSEYGTFCDNINSILNEVN<br>DLLDITQLQVANALMQGVTLSSNLNTNLHSDVDNIDFKSLLGCLGSQCGSSSRSLLEDLLFNKVKLSDVGFVEAYNNC<br>TGGSEIRDLLCVQSFNGIKVLPPILSETQISGYTTAATVAAMFPPWSAAAGVPFSLNVQYRINGLGVTMDVLNKNQKL<br>IANAFNKALLSIQNGFTATNSALAKIQSVVNANAQALNSLLQQLFNKFGAISSSLQEILSRLDNLEAQVQIDRLINGR<br>LTALNAYVSQQLSDITLIKAGASRAIEKVNECVKSQSPRINFCGNGNHILSLVQNAPYGLLFIHFSYKPTSFKTVLVS<br>PGLCLSGDRGIAPKQGYFIKQNDSWMFTGSSYYYPEPISDKNVVFMNSCSVNFTKAPFIYLNNSIPNLSDFEAELSLW<br>FKNHTSIAPNLTFNSHINATFLDLLIKRMKQIEDKIEEIESKQKKIENEIARIKKIKLVPRGSLEWSHPQFEK |

**Table S3. Experimental parameters for confocal microscopy.**

| Fig. | z-stack | pinhole size | objective lens | dichroic mirror + filters | image size |
|---|---|---|---|---|---|
| 4a | 24 × 1 μm | 31.93 μm | Plan Fluor 40× Oil DIC | 405/488/543/640<br>525/50<br>595/50 | 512×512 |
| 4c | 89 × 0.15 μm | 67.69 μm | Apo 100× Oil DIC | 405/488/561<br>525/50<br>595/50 | 1024×1024 |
| 4d | 30 × 1 μm | 28.1 μm | Plan Fluor 40× Oil DIC | 405/488/561<br>525/50<br>595/50 | 512×512 |
| 4e | 52 × 0.41 μm | 45.98 μm | Apo 60× Oil λS DIC | 405/488/543/640<br>525/50<br>595/50 | 1024×1024 |
| S19b, c | 30 × 1 μm | 28.1 μm | Plan Fluor 40× Oil DIC | 405/488/561<br>525/50<br>595/50 | 512×512 |
| S20 | 41 × 0.5 μm | 35.76 μm | Apo 60× Oil λS DIC | 405/488/561<br>525/50<br>595/50 | 512×512 |
| S21 | 55 × 0.41 μm | 45.98 μm | Apo 60× Oil λS DIC | 405/488/543/640<br>525/50<br>595/50 | 1024×1024 |
| S22 | 30 × 1 μm | 28.1 μm | Plan Fluor 40× Oil DIC | 405/488/561<br>525/50<br>595/50 | 512×512 |
